# Ceramic Packaging in Neural Implants

**DOI:** 10.1101/2020.06.26.174144

**Authors:** Konlin Shen, Michel M. Maharbiz

## Abstract

The lifetime of neural implants is strongly dependent on packaging due to the aqueous and biochemically aggressive nature of the body. Over the last decade, there has been a drive towards neuromodulatory implants which are wireless and approaching millimeter-scales with increasing electrode count. A so-far unrealized goal for these new types of devices is an in-vivo lifetime comparable to a sizable fraction of a healthy patient’s lifetime (>10-20 years). Existing, approved medical implants commonly encapsulate components in metal enclosures (e.g. titanium) with brazed ceramic inserts for electrode feedthrough. It is unclear how amenable the traditional approach is to the simultaneous goals of miniaturization, increased channel count, and wireless communication. Ceramic materials have also played a significant role in traditional medical implants due to their dielectric properties, corrosion resistance, biocompatibility, and high strength, but are not as commonly used for housing materials due to their brittleness and the difficulty they present in creating complex housing geometries. However, thin film technology has opened new opportunities for ceramics processing. Thin films derived largely from the semiconductor industry can be deposited and patterned in new ways, have conductivities which can be altered during manufacturing to provide conductors as well as insulators, and can be used to fabricate flexible substrates. In this review, we give an overview of packaging for neural implants, with an emphasis on how ceramic materials have been utilized in medical device packaging, as well as how ceramic thin film micromachining and processing may be further developed to create truly reliable, miniaturized, neural implants

## 1. Introduction

As is well known, the ability to interface with nervous tissue has great therapeutic potential, promising to treat a wide range of conditions ranging from motor dysfunction (Parkinsonian tremor, tetraplegia, Meige syndrome, etc.), psychiatric disorders (depression, OCD, etc.) (Chen, et al., 2017, Sun & Morrell, 2014), and peripheral nerve applications (Tracey, 2002, Borovikova, et al., 2000, Bosch & Groen, 2000, Slavin, et al., 2006) among others. Additionally, bidirectional communication with the nervous system allows for continuous monitoring and modulation of the nervous system, such that preventative actions can be taken to slow or even halt the progression of neuropathies (Famm, et al., 2013, Rouse, et al., 2011, Poon, 2014). Commercial efforts to deploy these types of interventions through the use of active implantable medical devices (AIMDs) is growing rapidly; an oft-cited estimate shows that the neuromodulation device industry was valued at 8.4 billion dollars in 2018 and is expected to grow to 13.3 billion dollars by 2022 (Cavuoto, 2018).

### 1.1 New and Existing Challenges in clinical neural implant packaging

The majority of neuromodulation devices rely on metal electrodes. Most of the electrochemical action occurs at the electrode-tissue interface; electrical current injected by the device changes the chemical potential of the local environment of a nerve, modulating the activity of nerve targets to achieve some therapeutic effect. These electrodes can also be used to detect extracellular changes in chemical potential, which can be used as a proxy for neuronal or peripheral nerve activity. A typical neural implant consists of: 1) microelectronics which assist in recording neural signals from or transmitting stimulation waveforms to metal electrodes; 2) the metal electrodes which interface and make contact with tissue; 3) insulated, conductive leads which connect the electrodes to the microelectronics; and 4) a battery, antenna coil, wires and/or other means to provide power to the implant. Depending on the application, emerging implants may need to interface with hundreds or thousands of electrode channels (Cha, et al., 1992, Lovell, et al., 2005, Buzsáki, 2004, Viventi, et al., 2011, Chang, 2015, Chung, et al., 2019). For most applications, the implantation is expected to be performed once (i.e. no explantation is needed) and would remain functional for the patient’s lifetime. Viewed from the perspective of implant mechanical design and material choices, both the tissue reaction to the implant (biotic) and the implant’s response to the tissue environment (abiotic) must be considered to achieve a long lifetime in-vivo (Fig. 1). Percutaneous connections can serve as routes for infection, so ideally all four components of the neural implant are fully implanted within the body.

**Fig. 1.**
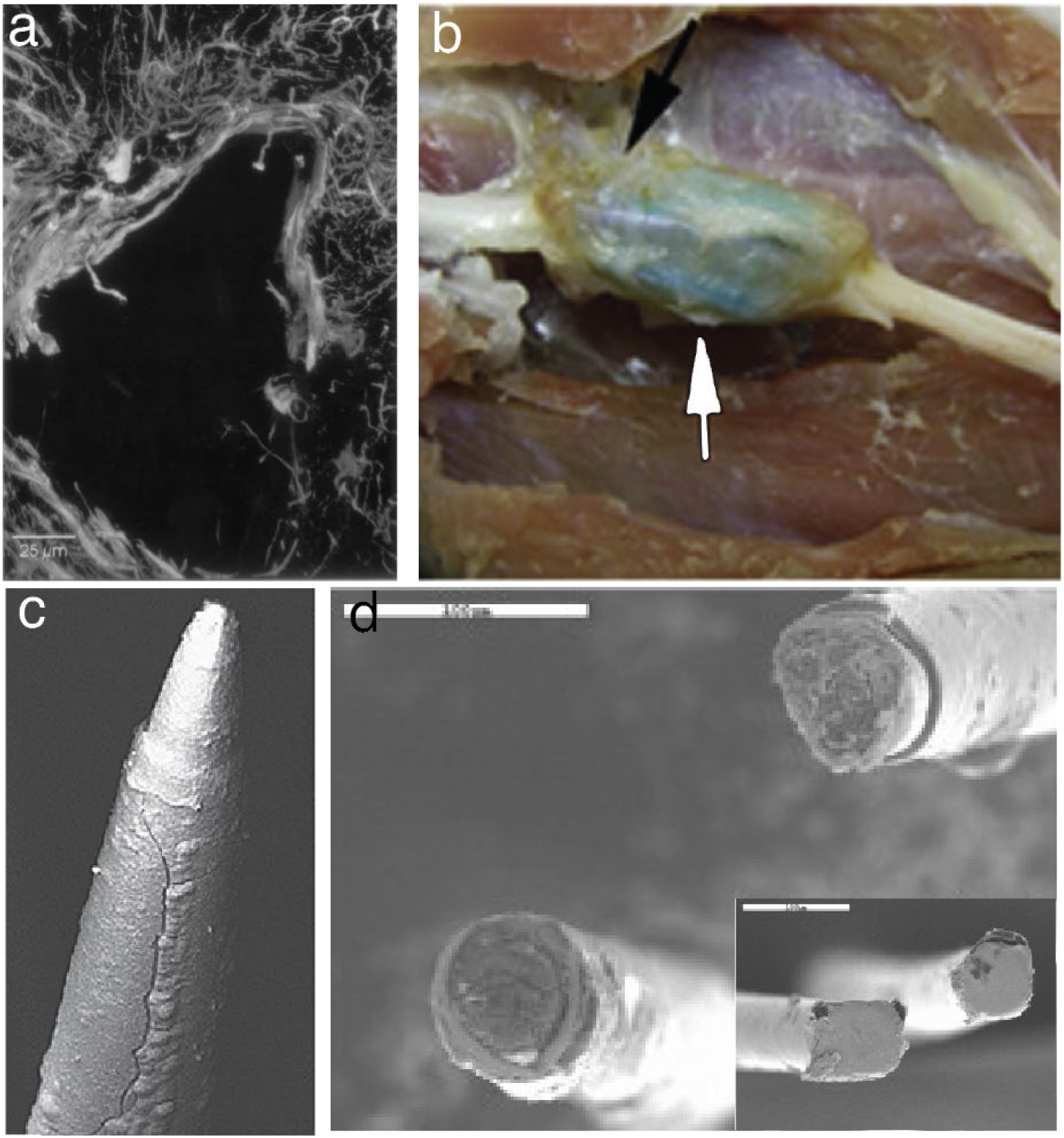
Failure modes of neural implants. a) Stain for glial fibrillary acidic protein (GFAP) in rats after explanting a penetrating neural probe left in the cortex for 12 weeks. The formation of a thick glial sheath around the implant site is clear, separating the implant from neurons (Turner, et al., 1999). b) Formation of a fibrotic capsule in the cat sciatic nerve after 5-6 weeks post implantation of a microelectrode array. The arrows indicate where the device was implanted (Christensen, et al., 2014). c) Cracks in the parylene encapsulation of a microelectrode array after 554-days post implantation in the cortex of a non-human primate (Barrese, et al., 2013). d) Corrosion in tungsten microwires after 87 days post-implantation in rat cortex. Inset shows the microwires prior to implantation. Roughening of the microwire surface, as well as delamination of the polyimide encapsulation can be seen (Patrick, et al., 2011).

Once the device is implanted, the body mounts an immune response against the device. This has been well-studied and covered by numerous reviews (Kozai, et al., 2015, Polikov, et al., 2005, Potter, et al., 2012, Szarowski, et al., 2003). A hallmark event of the immune response to implanted foreign body is gliosis in the central nervous system (Wellman, et al., 2019, Seymour & Kipke, 2007, Lind, et al., 2013, Turner, et al., 1999) and fibrosis in the peripheral nervous system (Romero, et al., 2001, Grill & Mortimer, 2000, Christensen, et al., 2014). While fibrosis can occasionally be useful to stabilize implants, the presence of scar tissue in both the central nervous system and peripheral nervous system reduces the efficacy of implanted devices by walling off healthy tissue from the electrodes (Fig. 1a,b). However, smaller implants have been found to have reduce scar tissue encapsulation around the device (Seymour & Kipke, 2007). Furthermore, large implants require more invasive surgeries and can also result in discomfort for the patient, or worse, skin erosion (Peña, et al., 2008). Thus, miniaturizing implants is highly desirable to increase the efficacy of chronic implants.

In parallel, the effect of the tissue environment on the implant material stack is also critical. The tissue environment is aqueous, saline, and both chemically rich and chemically aggressive. The immune response mounted by the body in response to an implant releases reactive oxidative species (ROS) which attack and degrade the implant (Fig. 1c) (Patrick, et al., 2011, Takmakov, et al., 2015). Corrosion of exposed metals is a major concern, with mitigation strategies heavily relying on material choices (Fig. 1d). Due to the saline environment of the tissue, galvanic corrosion between exposed metals can occur both in dissimilar metals and between identical metals in different environments (crevice corrosion) (Gilbert, et al., 1993, Kennell & Evitts, 2009). Corrosion can also be exacerbated by electrical stimulation due to electrolysis, which reduces the pH around the implant (Donaldson & Sayer, 1981, James, et al., 2016).

The consequence of these abiotic failure modes is eventual loss of hermeticity. This can result in moisture entering the packaging, shorting and corroding microelectronics. Simultaneously, non-biocompatible materials from the packaged microelectronics may leach into tissue, causing cell death and/or exacerbating the foreign body response (Vanhoestenberghe & Donaldson, 2013). Additionally, the packaging must be a good molecular barrier beyond water vapor since diffusion of mobile ions such as Na^+^, K^+^, and Cl^−^ into the integrated circuits within the housing can cause premature failure of the implant (Song, et al., 2017).

Putting the aforementioned requirements together, the ideal implant is capable of high channel counts while being extremely miniaturized, wireless, biocompatible, biostable, and ultimately hermetic for timescales commensurate with a patient’s lifetime. The technology limitation is not in the microelectronics or the interface, as many miniaturized devices with hundreds of channels have been demonstrated (Zhou, et al., 2019, Jun, et al., 2017, Raducanu, et al., 2017, Viventi, et al., 2011, Stevenson & Kording, 2011), but rather in the packaging of the implant components.

### 1.2 Packaging Materials for Medical Implants

Previous iterations of AIMDs have generally relied on polymeric potting (i.e. insulation and mechanical protection) or metal housings. For example, the original pacemakers were encapsulated in thick epoxy before titanium housing became the standard housing for implants (although the leads between internal pulse generators and their electrodes are still potted in silicone). Many researchers have long seen the choice between polymeric encapsulation and metal housing as equivalent to using a conformal material vs the manufacturing of an impermeable envelope (Donaldson, 1976). Unfortunately, while polymer encapsulation materials (including silicone, parylene, polyimide, and epoxy) have the advantage of being highly conformal, inexpensive, and easy to apply, polymeric materials are not good long lifetime materials due to their relatively high water vapor permeabilities (Traeger, 1977). This is often circumvented for larger implants by increasing polymer thickness to many millimeters (or more) but it is not a viable strategy for very thin layers in miniaturized devices. In addition, the ROS species discussed earlier are well-known to participate in the breakdown of polymer coatings. For example, parylene is known to crack after months of implantation in the body (Barrese, et al., 2013, Takmakov, et al., 2015). Additionally, polymers such as parylene and liquid crystal polymers will undergo material property changes at high temperatures, which prevents the use of steam autoclaves, which generally operate at 120 °C. While ethylene oxide (EtO) sterilization can be used at much lower temperatures, the permeability of polymeric potting materials can allow both EtO and moisture to be stored in the material bulk and a degassing step is required (Stieglitz, 2010).

Due to their very low water vapor permeabilities, metals have been the industry standard for housing implantable microelectronics. In particular, titanium has been the material of choice for AIMD housings due to the formation of a stable and inert oxide passivation layer (Bowman & Meindl, 1986). While titanium housings have been an excellent workhorse during the past 50+ years, they have some limitations. For example, metals alone cannot be used in packaging, as the electrical feedthroughs which connect the electrodes to the microelectronics must be electrically isolated from each other. More conspicuously, the use of metal-only housings generally precludes the ability to efficiently transfer energy wirelessly using electromagnetics without undue heating of the package caused by eddy currents.

In contrast to metals, ceramic materials not only have low water vapor permeabilities, are transparent to RF, can be rendered optically transparent, and, recently, have been shown to be amenable to ultrasonic power transfer (Gutierrez, et al., 2017). Many ceramics are also chemically inert, biocompatible, electrically insulating, and harder and stronger than most metals (Piconi & Maccauro, 1999, Vlasov & Karabanova, 1993). Due to these favorable properties, ceramics have had a long history as implant materials both alone and in conjunction with titanium housing. As implants become more ubiquitous and continue to shrink, ceramics will play an increasingly important role in the development of active implantable medical devices.

### 2. Classical ceramic packaging methods

Currently, the major application of ceramics in AIMDs is in hermetic feedthroughs, which provide pathways for electrical signals to be routed into and out of the implant housing. The development of the alumina-titanium feedthrough for the cardiac pacemaker significantly increased the longevity of AIMDs from months to years and the hermetic alumina-titanium paradigm has been a mainstay of the AIMD industry ever since (Ruys, 2018). Ceramics have also been used to improve wireless transmission in titanium housings, acting as windows with some degree of RF transparency (Borton, et al., 2013, Bjune, et al., 2015). As a clinical example, the Axonics sacral nerve stimulator utilizes a hybrid titanium-ceramic package for the implantable pulse generator (IPG) which allows for high efficiency electromagnetic coupling (Elterman, 2018). This unique construction allows for an extremely small charging coil, enabling the Axonics IPG to be as small as 5 cm^3^. In contrast, other IPGs housed solely in titanium are on the order of tens of cm^3^. While completely ceramic housings have been explored, particularly in cochlear implants in which wireless communication is essential, the brittleness of ceramic has been a reliability concern. In cochlear implants, all-alumina housing was explored by Cochlear Ltd, MED-EL, and Neurolec, but due to the large amount of exposed area, the housings were vulnerable to failure due to mechanical impact (Fig. 2a) (Stöver & Lenarz, 2009). Similarly, the first generation BION microstimulator used a glass housing to enhance RF transparency, but found that the housing was prone to fracture after repeated bending stress due to muscle contraction (Fig. 2b) (Loeb, et al., 2007). Careful mechanical design was able to improve the strength of the BION glass housing; subsequent versions of the implant utilized a ceramic housing with a 7x improvement in bending strength (Kane, et al., 2011). Arguably, the reason glass worked at all for the BION was the small size and relatively thick glass walls. For larger implants, titanium housings remain the preferred solution. Thus, while ceramics may not comprise the majority of modern implant packages, they are an essential component. To this end, whether the ceramic part acts as a communication window, substrate, or feedthrough, it must be shaped and then joined to the rest of the implant.

**Fig. 2.**
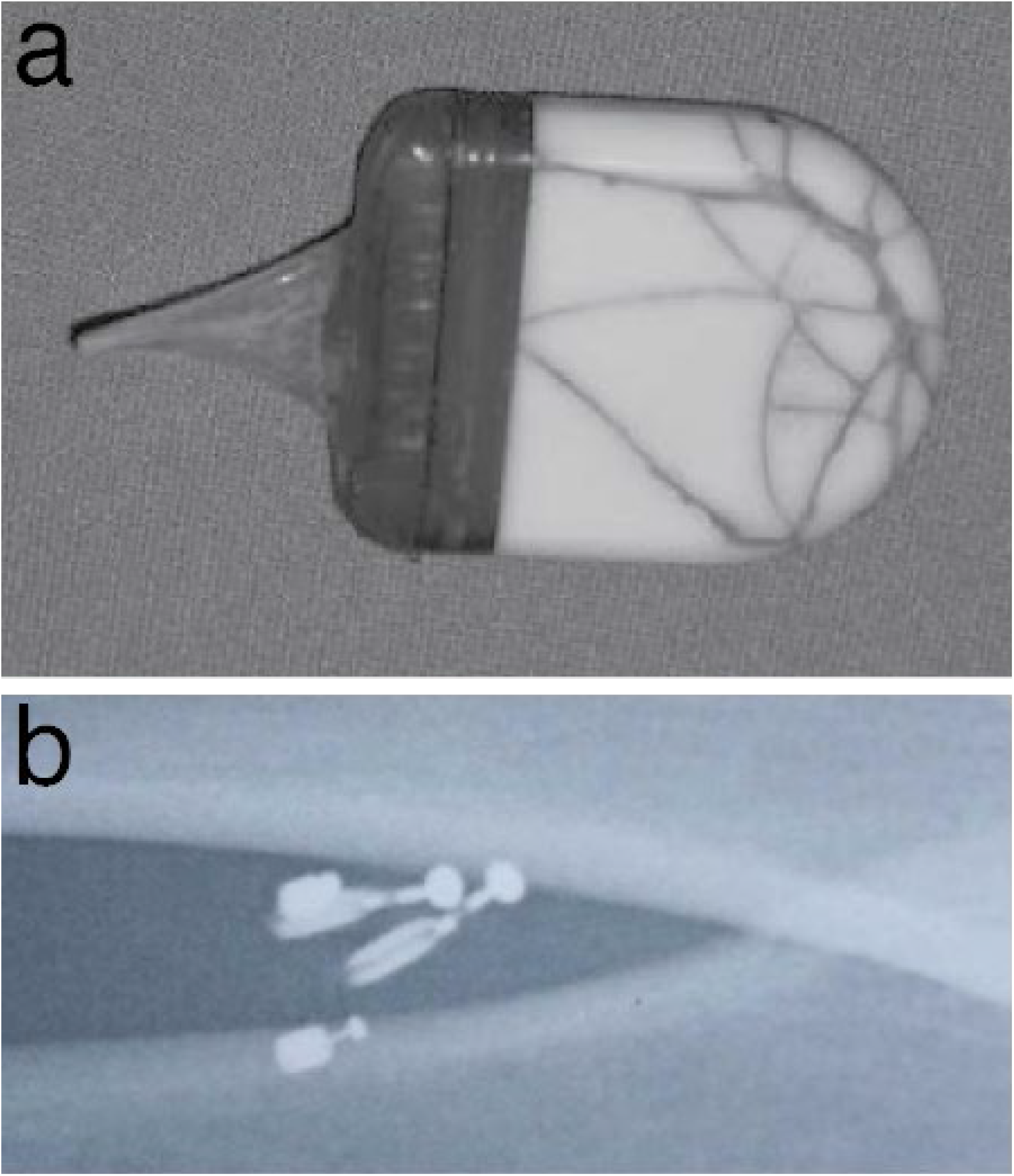
Mechanical failure of ceramic housings. a) Cracked alumina casing in a cochlear implant due to mechanical impact of the head (Stöver & Lenarz, 2009). b) Broken intramuscular microstimulators encased in glass housing due to bending stresses from muscle contractions (Loeb, et al., 2007).

### 2.1 Forming of ceramic packaging components

Conventionally, ceramic parts are created from powder, which is consolidated and shaped into a “green body”, then fired to densify the final part (Fig. 3). Shaping of the green body can be done in a variety of ways but is dominated by “wet-forming” and “dry-forming” methods. In wet-forming methods, the ceramic powder is added to a liquid medium to form a slurry in which binders and plasticizers can be added. This slurry can then be poured into a mold (injection molding) to form a complex shape; tapecast into a sheet for green machining; or slipcast to form a hollow shape; among other methods (Yeong, et al., 2013). In dry-forming methods, the ceramic powder is simply pressed to form the desired shape using methods like uniaxial die pressing or cold isostatic pressing (Ruys, 2018). When forming ceramics, the major concern is avoiding structural defects, such as cracks. These cracks tend to occur during drying processes, such as the drying step needed to remove organic binders and solvents from ceramic slurries. Dry-forming does reduce the risk of cracking, but particle packing homogeneity and shape complexity are worse than for wet-forming methods. Injection molding allows for complex parts with high precision and have been used to form the “comb” style alumina feedthroughs used in cochlear implants (Fig. 4b). However, injection molding can be limited; particularly complex geometries may result in short shots (incomplete filling of the mold) or knit-lines (improper mixing of the slurry from multiple injection points resulting in stress-lines) (Christian & Kenis, 2007). The main method of forming ceramic substrates, particularly for electronics, has been tape-casting; this method can create ceramic substrates with sub-mm thickness (Tummala, 1991). With this method, a ceramic slurry is doctor-bladed across a surface with a controlled velocity and height to achieve a green sheet with a target thickness. This sheet can then be machined and fired to form the final substrate. 3D printed ceramics are also a promising new technology, allowing for very rapid and cheap manufacturing since they do not require tooling. However, in its current state, 3D printing cannot achieve the same thin substrates as tapecasting; additionally, dimensional accuracy, surface finish and high porosity remain issues (Travitzky, et al., 2014). As in many material domains, the prospect of being able to rapidly and cheaply manufacture ceramic parts with high geometric complexity is accelerating the development of 3D printed ceramics; with improvements, additive manufacturing may soon replace techniques like injection molding and cold isostatic pressing (Chen, et al., 2019).

**Fig. 3.**
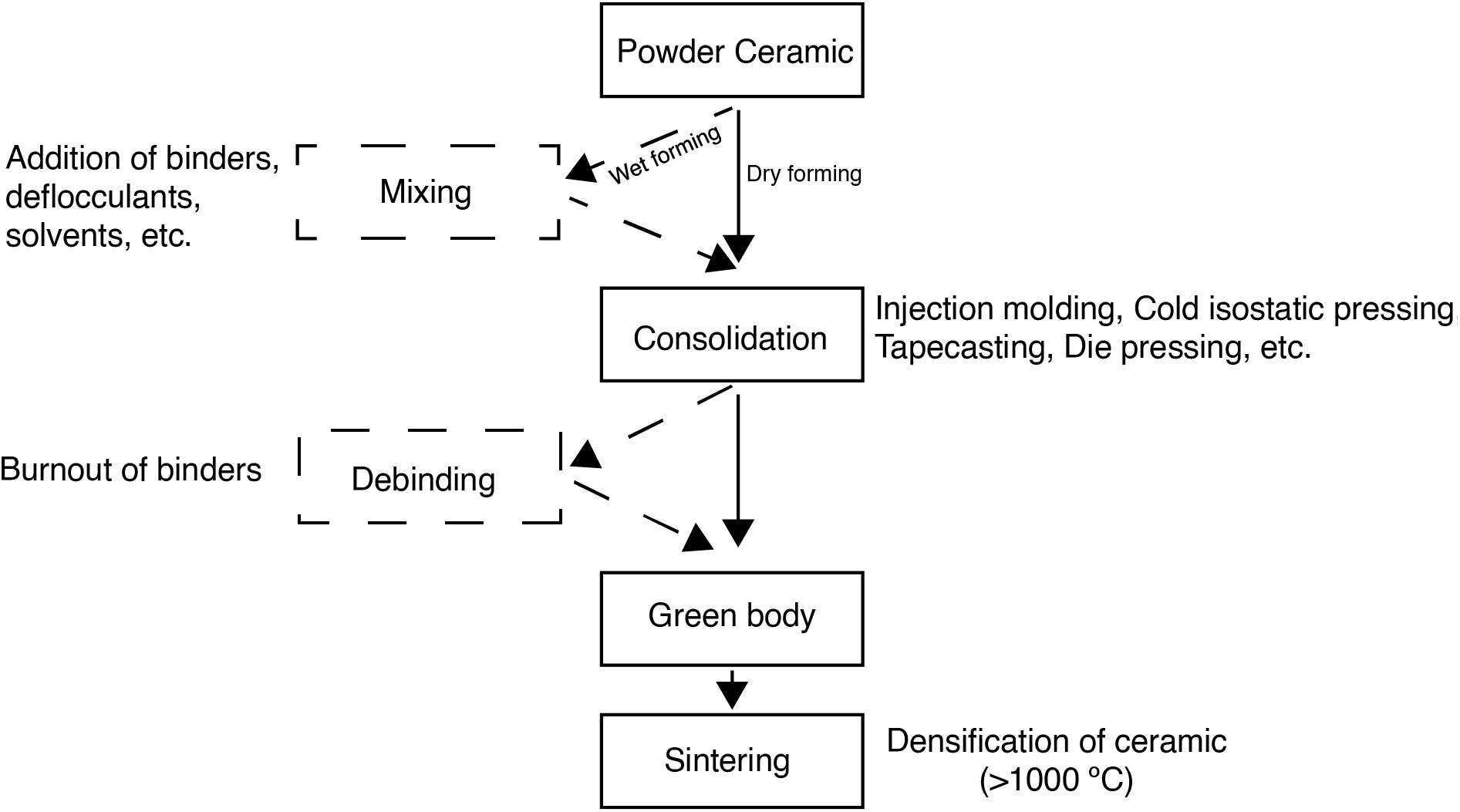
Flowchart of the conventional ceramic forming process. Powder ceramic powder is shaped during the consolidation step, with the method of consolidation depending on whether the powder has been mixed into a slurry (wet forming) or not (dry forming). Post consolidation, binders are burned out to form a green body. The green body is then sintered, in which the ceramic particles enlarge and join together to form a dense final product.

**Fig. 4.**
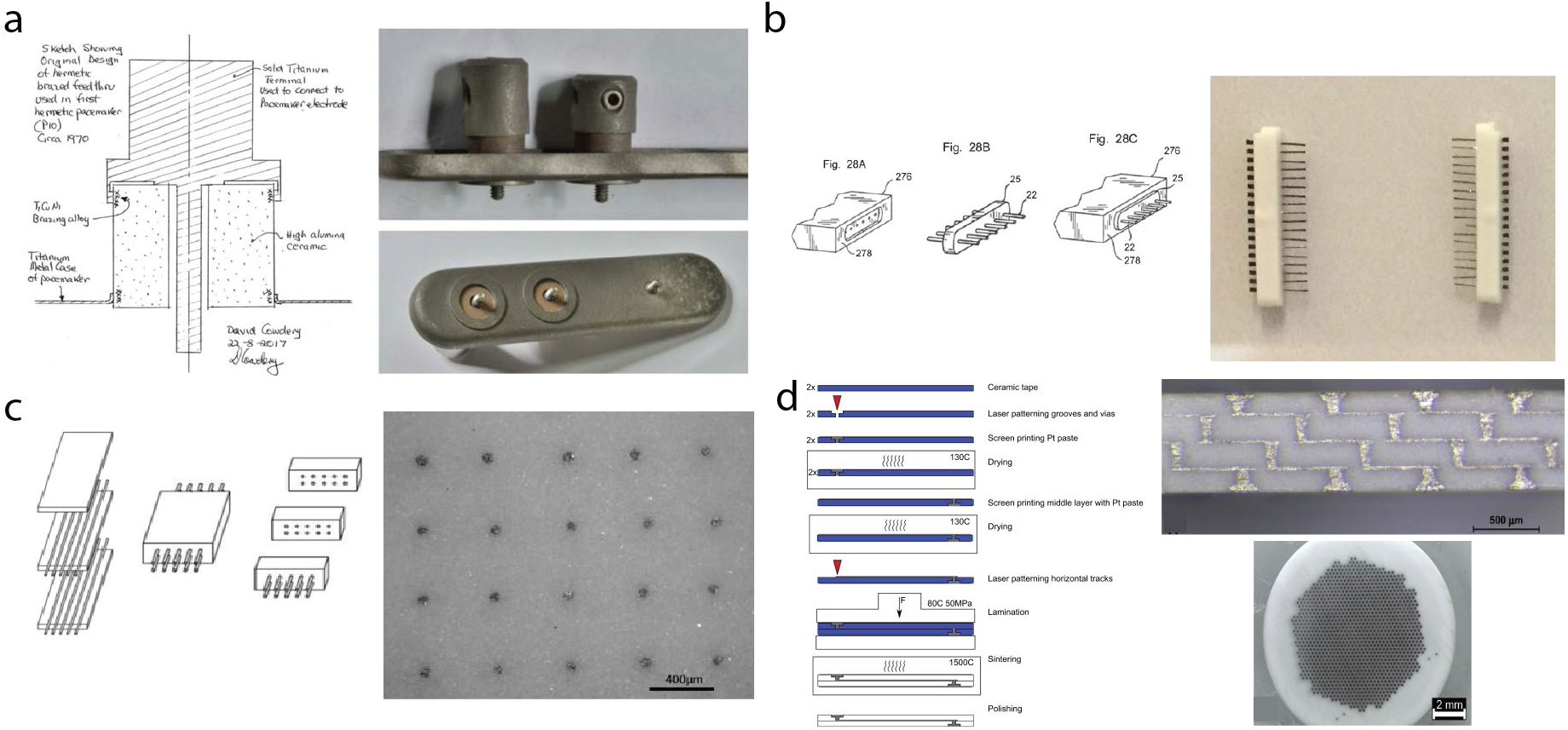
Traditional ceramic forming methods. a) The original Alumina/Ti feedthrough. Here, an alumina feedthrough is brazed to a titanium housing, with a titanium pin brazed inside it. (Ruys, 2018) b) The “comb” feedthrough using Pt wires, brazed into alumina feedthroughs which are then brazed to a Ti package using the same procedure as a) (Ruys, 2018). c) Co-fired Pt-Alumina feedthroughs. Pt wires are laid over Alumina green body “loaves”, then pressed together and fired. The sintered ceramic parts are then diced and ground to produce an array of Pt feedthroughs (Gill, et al., 2013). d) Co-fired Pt-alumina feedthroughs using horizontal interconnects and Pt paste. Alumina green bodies are laser patterned and Pt paste is screen printed to fill vias and interconnects in multiple layers. The layers are then laminated together and sintered as one piece. This method can yield an extremely high density of Pt interconnects in alumina substrates and the use of the serpentine design increases the leakage path to improve hermeticity (Green, et al., 2013).

### 2.2 Joining and metallization of ceramics

Once the ceramic part is formed, it must be hermetically joined to the housing material to form feedthroughs, windows or seals. In a feedthrough, one or more metal pins are also joined to the ceramic body to create conductive paths between the outside and the inside of the package; the ceramic body is then joined to the metal housing of the implant (Fig. 4a). The most common method of joining ceramic is brazing, in which two materials are joined with a filler material at a temperature exceeding 450 °C (differentiating it from soldering, which occurs at lower temperatures). Brazing fillers can be either passive or active. In an active braze alloy, the paste generally contains an active element such as Ti or Zr which promotes the adhesion and wetting of the ceramic material (Apollo, et al., 2016). As such, brazing with active braze alloys (ABA) generally requires a vacuum furnace, capable of pumping down to at least 10^−5^ torr. ABAs are widely used hermetic joining of feedthroughs and housing sealing. A classic example is the Ti-Cu-Ni brazing alloy used in alumina-titanium bonding process; the Ti content promotes wetting of the braze alloy on the alumina, and endows the joint with high biocompatibility and corrosion resistance. Partially because of objections to the cytotoxicity of copper, copper-free braze alloys have been explored. Ti-Ni brazes have been shown to hermetically seal zirconia-to-titanium (Jiang, et al., 2005) and pure Ni itself was found to be able to hermetically seal the same materials because the titanium from the part was able to diffuse into the nickel to form a eutectic alloy (Jiang & Zhou, 2009). Both gold and silver ABAs have been utilized to hermetically seal polycrystalline diamond (PCD) packages; the high gold content of the gold-ABA arguably promotes biocompatibility (Lichter, et al., 2015). In the last example, it was found that the gold-ABA did not wet the PCD well, but the silver-ABA could be used as an intermediate layer, with the silver layer not exposed to the tissue environment after encapsulation by the gold layer.

In contrast, passive braze alloys do not contain reactive elements to promote wetting, which allows them to be used in low vacuum environments. Typically, these braze materials are noble metals such as gold or platinum (Siddiqui & Jones, 2014). Unfortunately, passive alloy joints are generally mechanically weaker and somewhat porous (Agathopoulos, et al., 2002, Correia, et al., 1998). Furthermore, the poor wetting of these alloys to ceramic materials requires a metal interlayer. This can be achieved through physical vapor deposition, chemical vapor deposition, screen printing, or other methods (Santella & Pak, 1993).

While hermetic brazed feedthroughs have had much success, they are limited in pin (conductor) density due to the risk of the ceramic cracking as the conductors are brought closer together (Kelly, et al., 2011). The density limit can be increased integrating metal parts or metal pastes with the ceramic green body and simultaneous firing, a method also known as co-firing (Schuettler & Stieglitz, 2013). Platinum is used for this due to its high melting temperature, low resistivity, and ability to form solid-state bonds with alumina (Allen & Borbidge, 1983, Guenther, et al., 2014). Solid Pt wires can be cofired with the ceramic, in which wires are placed between “loaves” of green sheet, compressed together, and fired (Fig. 4c) (Gill, et al., 2013). This method has been able to create 10 x 10 arrays of feedthroughs with pitches between 400 and 450 μm. When using metal pastes, slurries containing metal particles are screen printed into vias machined into the green body and sintered along with the ceramic (Fig. 4d) (Yang, et al., 2018, Green, et al., 2013). Afterwards, the ceramic substrate with metal vias can be metallized again to form interconnects. Substrates with ~1000 feedthroughs and pitches on the order of 40 μm have been demonstrated. To date, no other method has been able to provide such high feedthrough densities.

One drawback of brazing and co-firing is the high temperature required. During high temperature excursions, thermal expansion coefficient mismatches between different parts can result in cracking or loss of hermeticity. Furthermore, due to the use of organic binders in metal paste feedthroughs, sintering of these pastes can result in void formation which also compromises the hermeticity of the via. Finally, where hermetic sealing of the housing is concerned, high temperature joining is often incompatible with the microelectronic components within the housing. Thus, low temperature methods of joining ceramic are highly desirable.

A relatively new method for low temperature feedthrough formation is to form extruded metal vias with stud bumping (Fig. 5) (Shah, et al., 2012, Langenmair, et al., 2018). In this method, vias are machined into a ceramic substrate and subsequently metallized. Gold bumps are placed over the vias using an ultrasonic ball bonder to alloy the gold with the via metallization, then coined. In this way, extremely dense hermetic feedthroughs in a ceramic substrate can be made at a low temperature, with a theoretical density up to 2500 feedthroughs/cm^2^. While the feedthroughs themselves have been found to be hermetic, to date, substrates with over 100 feedthroughs have been found to be non-hermetic due to misalignment between the gold stud and the via. In published attempts, studs were manually bonded, which contributed to misalignment error. Further work illustrating that automation of the stud placement to improve alignment also improves hermeticity is needed to validate this technique.

**Fig. 5.**
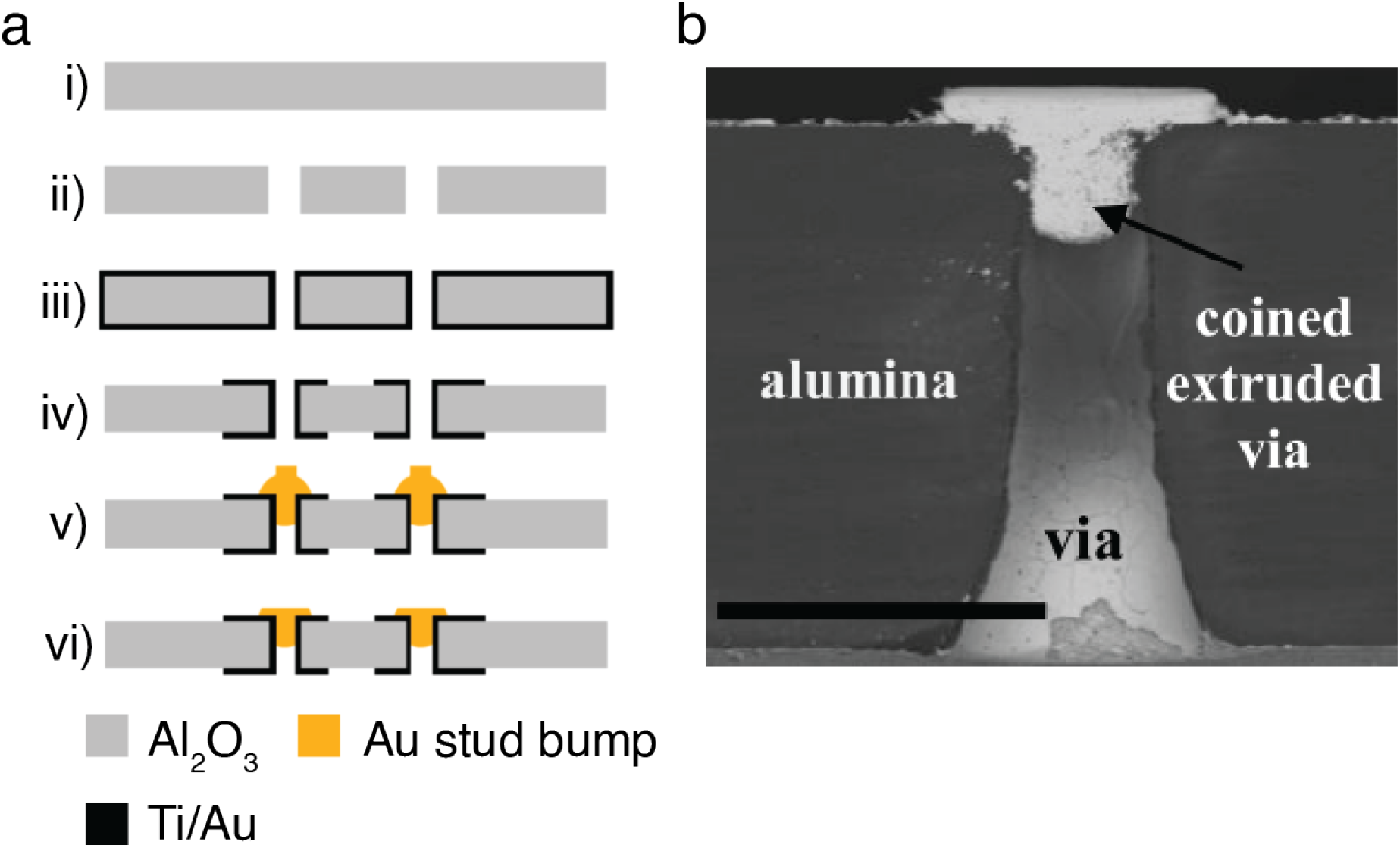
Extruded metal vias concept. a) Process flow for creating extruded metal vias in ceramic. i) a ceramic (alumina) substrate ii) vias are created by laser machining iii) Ti/Au metallization is conformally sputtered into vias for sidewall coverage iv) metallization is lithographically patterned on both sides v) Au stud bumps are positioned and extruded over the vias using an ultrasonic ball bonder vi) Au stud bumps are then coined using a flip chip bonder to planarize the substrate and improve hermiticity. b) cross sectional view of an extruded via (scale bar = 100 μm) (Shah, et al., 2012)

Other approaches for sealing packages at low temperature have been explored. Schuettler et al. developed a 360-channel vision prosthetic on a ceramic substrate sealed to a metal cup with low temperature solder in a custom hermetic sealing chamber (Schuettler, et al., 2010, Schuettler, et al., 2010). While this method yielded devices with excellent hermeticity, the use of solder and the requirement of a solderable housing material left biocompatibility an open question. That work utilized a brass cup for solderability, as titanium is essentially impossible to solder (due to its surface oxide) but brass is cytotoxic as a copper-zinc alloy.

The standard practice for low temperature sealing of titanium housings remains laser welding (Xiansheng, et al., 2011, Gedopt & Delarbre, 2000). Laser welding allows for very localized deposition of heat to melt and fuse materials together. This is advantageous for sealing housing without thermal damage to internal components. However, local heating is problematic for joining ceramics due to thermomechanical stresses generated by the large temperature gradient between the laser affected zone (LAZ) and the areas outside of it (Witte, et al., 2002). Early efforts to circumvent this issue have been to reduce the gradient by preheating the part prior to welding with an oven or a secondary laser (De Paris, et al., 1991, Riviere, et al., 1994, Exner & Nagel, 1999). Alternative methods to utilize laser welding include the use of a filler material such as glass frit to aid joining (Wu, et al., 2010); this enables a form of “laser brazing” proposed for hermetic vision prostheses as well as to seal the previously described PCD implants (Vanhoestenberghe, et al., 2008, Lichter, et al., 2015). More recently, ultrafast laser welding has been introduced as a method for crack-free ceramic joining without needing to preheat the parts (Penilla, et al., 2019, Watanabe, et al., 2006, Zimmermann, et al., 2013). An in-depth explanation behind the interaction between ultrashort laser pulses and matter can be found in both Miyamoto et al. and Penilla et al. (Miyamoto, et al., 2014, Penilla, et al., 2019). Briefly, ultrashort laser pulses stimulate non-linear absorption processes in material and can result in embedded molten pools within the bulk material. Because of the short pulse time, the material outside the molten pool remains its original temperature. The embedded molten pool then acts as an elastic body and thus does not generate thermomechanical stresses during cooling. Ultrafast laser welding for ceramic joining is a significant milestone for enabling all-ceramic housing and promises hermetic seals formed without heat-damage to the packaged components. Ultrafast laser welding does have material constraints, but careful engineering or tailoring of the ceramic microstructure can circumvent these issues (Penilla, et al., 2019).

### 2.3 Hermiticity Testing

Ultimately, a package must protect the internal microelectronics from the body environment and prevent compounds from the microelectronics components from leaking into the body environment. Thus, after forming and joining, the package must be leak tested to ensure hermiticity. Hermeticity testing generally occurs in two phases – gross testing and fine testing. A gross leak is defined by the military standard MIL-STD-883 as a package with a standard air equivalent leak rate greater than 10^−5^ atm-cc/sec (Greenhouse, 2000). A common gross-leak test is the *bubble test*, in which a detector liquid with a low boiling point is forced into the package by ‘bombing.’ Bombing is injection of the detector medium into the package by immersing the sealed package in the detector and pressurizing the fluid to drive it into the package. Post-bombing, the package is placed in an indicator liquid which has a much higher boiling temperature. The temperature of the indicator liquid is then raised above the boiling temperature of the detector liquid but below the boiling point of the indicator. This condition is maintained for at least 30 seconds. If bubbles are observed, the package is declared to be a “gross leaker”. Presuming the package passes the gross leak test, it is then tested for fine leaks. The current standard method of fine leak testing is to use helium (He) because of its small atomic size and rarity in the ambient environment. Two methods exist for helium leak testing: bombing and backfilling. In bombing, sealed packages are placed into a high pressure He environment, which drives He into the package. The bombed packages are then placed into a chamber with a mass spectrometer and the rate of helium leakage is measured. Alternatively, backfilling may be used in which packages are sealed in a helium environment and then transferred to a mass spectrometer to measure helium leakage. The leakage of He, of course, is not the same as the leakage of water vapor; to predict the water leakage rate, Graham’s Law of Effusion is used (Vanhoestenberghe & Donaldson, 2011):

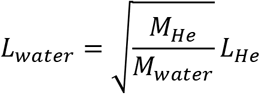

Where *L*_*water*_ and *L*_*He*_ are the leak rates for water and helium and *M*_*He*_ and *M*_*water*_ are molar masses for helium and water. For reasons which will be explored in the next section, it is difficult to pin down a He leak rate that guarantees hermiticity.

## 3. MEMS-enabled miniaturization of ceramic packages

In the pursuit of highly miniaturized implants, techniques from microelectromechanical systems (MEMS) and other semiconductor processing methods continue to be employed to create devices with sub-millimeter dimensions and to further increase electrode density (Seymour, et al., 2017). While the previously mentioned work by Green et al. demonstrated feedthrough densities on the order of 1000/cm^2^, it required a four-layer laminate structure, with each layer being roughly 200 μm thick (Green, et al., 2013). While the authors suggest that a two-layer structure is enough, this still results in a substrate at least 400 μm thick (not including the rest of the housing). Furthermore, the interconnects are structured using a laser machining process, which has a minimum line width of 20 μm. Microfabrication technology, however, can pattern sub-micron features on micron thick substrates and has a long history in the development of neural probes (Fig. 6) (Wise, 2005, Cheung, 2007). A representative, seminal, example is the technology developed at the University of Michigan. One of the early ‘Michigan Probes’ was only 20 μm wide, between 8 and 15 μm thick, between 1.5 and 3 mm long, and could support 10 channels (Fig. 6a) (Najafi, et al., 1985). This represents an electrode density of nearly 17000 channels/cm^2^! Modern neural probes have pushed this density even farther (Steinmetz, et al., 2018). The Neuropixel probe boasts 960 recording sites on a 1 cm long, 70 μm wide, and 20 μm thick silicon shank – an electrode density over 1 million channels/cm^2^ (Fig. 6b) (Jun, et al., 2017).

**Fig. 6.**
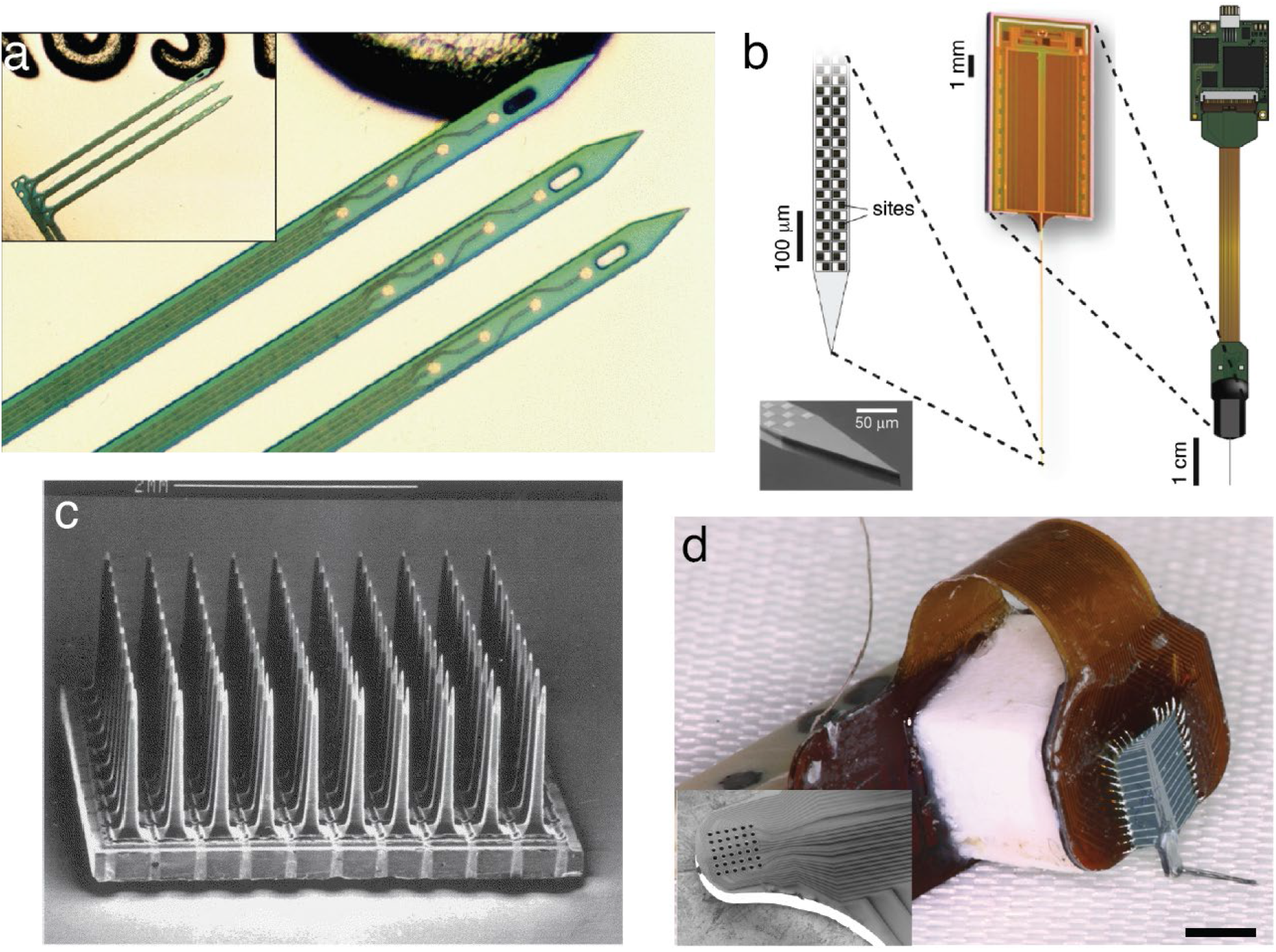
Examples of neural interfaces enabled by microfabrication technology. a) A Michigan probe. Here a 12-channel device is shown, with 4 channels on three separate shanks. The device is insulated by a combination of silicon dioxidie and silicon nitride. (Wise, 2005) b) The neuropixel probe. 960 recording sites are checkerboarded across a single shank, 70 μm wide x 1 cm long. This device has one of the highest electrode densities in neural recording devices (Steinmetz, et al., 2018, Jun, et al., 2017). c) The classic Utah array, a 10 x 10 penetrating microelectrode array developed at the University of Utah. The tines are made of doped silicon, singulated with a dicing saw, and chemically sharpened. The channels are all electrically isolated from each other using a “moat of glass” which surrounds the base of each electrode (Normann & Fernandez, 2016). d) A 32-channel penetrating carbon fiber microelectrode array in the style of the Utah array. The electrodes are 5 μm diameter carbon fibers, threaded through a silicon substrate. Inset: an SEM of the head of the Si substrate prior to carbon fiber insertion, which shows 20 μm vias with a center-to-center pitch of 38 μm (Massey, et al., 2019).

### 3.1 The challenge of miniaturized packages

Although microfabrication is indisputably a favored manufacturing route to small implants and high densities, severe issues arise, not all of them obvious. For example, the decreasing volumes of microfabricated housings result in a higher likelihood of reaching dew point, resulting in moisture condensation. Thus, smaller packages must have higher hermeticity constraints than larger packages. The relationship between package volume and maximum leak rate can be expressed by (Guenther, et al., 2020):

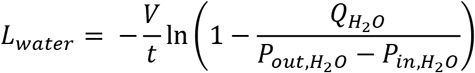

Where 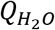 is the maximum allowed water partial pressure within the cavity, 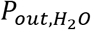 and 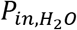 are the initial partial pressures of water on the outside and inside of the housing respectively, *V* is the free-volume of the housing, and *L*_*water*_ is the maximum leak rate allowed to maintain hermiticity within a time period *t*. The maximum allowed water partial pressure is usually referenced to the MIL-STD-883 recommended value of 5000 PPM, which is derived from the Clausius-Claperyon relationship as the maximum permissible water vapor pressure before condensation occurs.

This relationship between hermeticity (measured in He leak rate), housing free-volume, and moisture content is plotted in Fig. 7, which shows the water vapor partial pressure in PPM of a package of a given size and He leak-rate after 10 years. The manufacturing consequence of this phenomenon is that the maximum allowed leak rate for such small packages becomes too low to be detected by most He leak rate detectors currently employed, which are typically limited to 10^−11^ atm-cm^3^/s. Some methods have been used to try to improve the detection floor, such as cumulative helium leak detection (CHLD). In CHLD, a specialized cryo-pump is used which is capable of pumping out all other gases besides He from the test chamber, resulting in a higher detection sensitivity. However, Guenther et al. found several problems with this method when applied to microscale packages: outgassing of He from rubber o-rings created significant background noise, and the manufacturer’s requirement of having a test-chamber volume to sample-cavity volume ratio of less than 100 was very difficult without major modifications of the commercially available CHLD instruments (Guenther, et al., 2020).

**Fig. 7.**
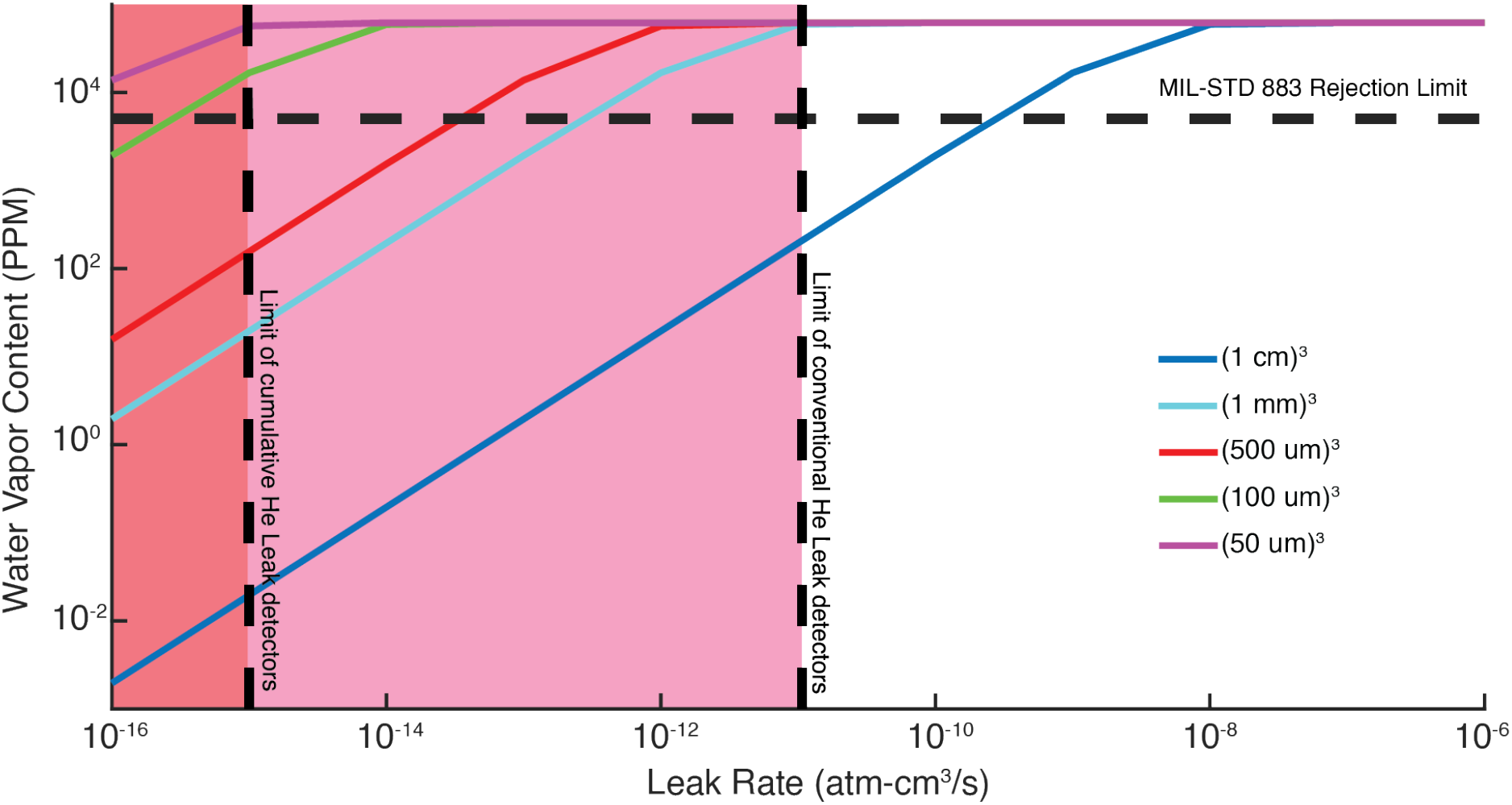
Predicted water vapor ingress after 10 years for a given leak rate and a given free volume. The maximum permissible water vapor content in a package is 5000 PPM as mandated by the MIL-STD-883 to prevent condensation. As volume decreases, the maximum allowable leak-rate to achieve a 10-year lifetime without reaching the maximum water vapor content decreases. Furthermore, the maximum leak-rate for small packages quickly becomes too low for conventional He leak rate detection and even cumulative He leak rate detection.

### 3.2 MEMS-enabled ceramic neural implants

Due to the difficulty of leak testing miniaturized housings, it follows that a potential path towards long-lasting hermetic implants is to conformally coat devices with packaging material, resulting in zero free-volume within the packaging. Thin film deposition techniques such as chemical vapor deposition (CVD) and atomic layer deposition (ALD) can result in highly conformal coatings to encapsulate implants. These thin films can be on the order of microns to sub-microns thick, which means that the packaging of the device will not add a significant amount of volume to the device. Furthermore, the ability to deposit a film over the microelectronics obviates the need for separate joining processes to seal housings. This is advantageous not only for miniaturization, but also in wireless transmission, where even the small amount of metal in the braze alloy can detune an antenna. Many ceramics can be deposited in this way and have been used as encapsulation materials. Again harkening back to the classic Michigan probe, the conductors on the probe were encapsulated by a bilayer of CVD silicon nitride (SiN) and silicon dioxide (SiO_2_) (Najafi, et al., 1985). Xie et al. demonstrated the use of ALD alumina (Al_2_O_3_) in combination with parylene as a hermetic bilayer, projecting a lifetime beyond 5 years at 37 °C (Xie, et al., 2013). Weiland et al. showed that 5 μm thick diamond-like carbon thin films were effective moisture and ion-barriers, showing only a single pinhole over a 1000 mm^2^ area and less than 1% change in electrical properties of an encapsulated integrated circuit (IC) (Weiland, et al., 2013).

The work done by Weiland et al. demonstrates another benefit of thin-film deposition technology: low temperature deposition. The high temperatures required for ceramic sintering and brazing for sealing housing are often incompatible with CMOS technology. However, thin-film deposition can often be done at much lower temperatures, with additional energy introduced to the system with plasma rather than heat. Amorphous silicon carbide (a-SiC) for example has been pursued as a coating for neural implants due to its biocompatibility and chemical inertness and can be deposited at temperatures lower than 400 °C through the use of plasma-enhanced CVD (PECVD) (Cogan, et al., 2003, Hsu, et al., 2007). Lei et al. demonstrated the use of PECVD a-SiC deposited at 350 °C as a dielectric coating for retinal implants in which 240 nm. SiC coated retinal prostheses were implanted in rats for up to 1 year, showing significantly less degradation than only SiN coated protheses (Lei, et al., 2016). ALD hafnium oxide (HfO_2_) coatings have also been explored. Jeong et al. deposited alternating layers of 10 nm thick HfO_2_ and SiO_2_ at 250 °C to form a 100 nm thick coating over an RF communication IC. The encapsulated ICs were demonstrated to work while submerged in 87 °C saline for over 180 days, suggesting much longer lifetimes at physiological temperatures (Jeong, et al., 2019).

Another major benefit of thin film ceramic coatings is low flexural rigidity which is a property that bulk ceramic does not have and limits its use as a housing material. As an example, microns-thick silicon carbide can be fabricated into compliant high channel count microelectrocorticography (ECoG) arrays flexible enough to be wrapped entirely around the sciatic nerve of a rat (Fig. 8a) (Diaz-Botia, et al., 2017). Similarly, methods for transferring sub-micron thick SiO_2_ layers over flexible electronics have been developed (Fang, et al., 2016). By laminating thermally-gown SiO_2_, Fang et al. demonstrated a 0.95 cm × 1.15 cm, 900-nm thick, 396 channel, capacitive sensor for electrophysiological measurement, which was flexible enough to fit to the curvature of cardiac tissue (Fig. 8b, d) (Fang, et al., 2017). That work has been extended to SiN, SiC, and HfO_2_ (Fig. 8c) (Song, et al., 2018, Song, et al., 2017, Phan, et al., 2019).

**Fig. 8.**
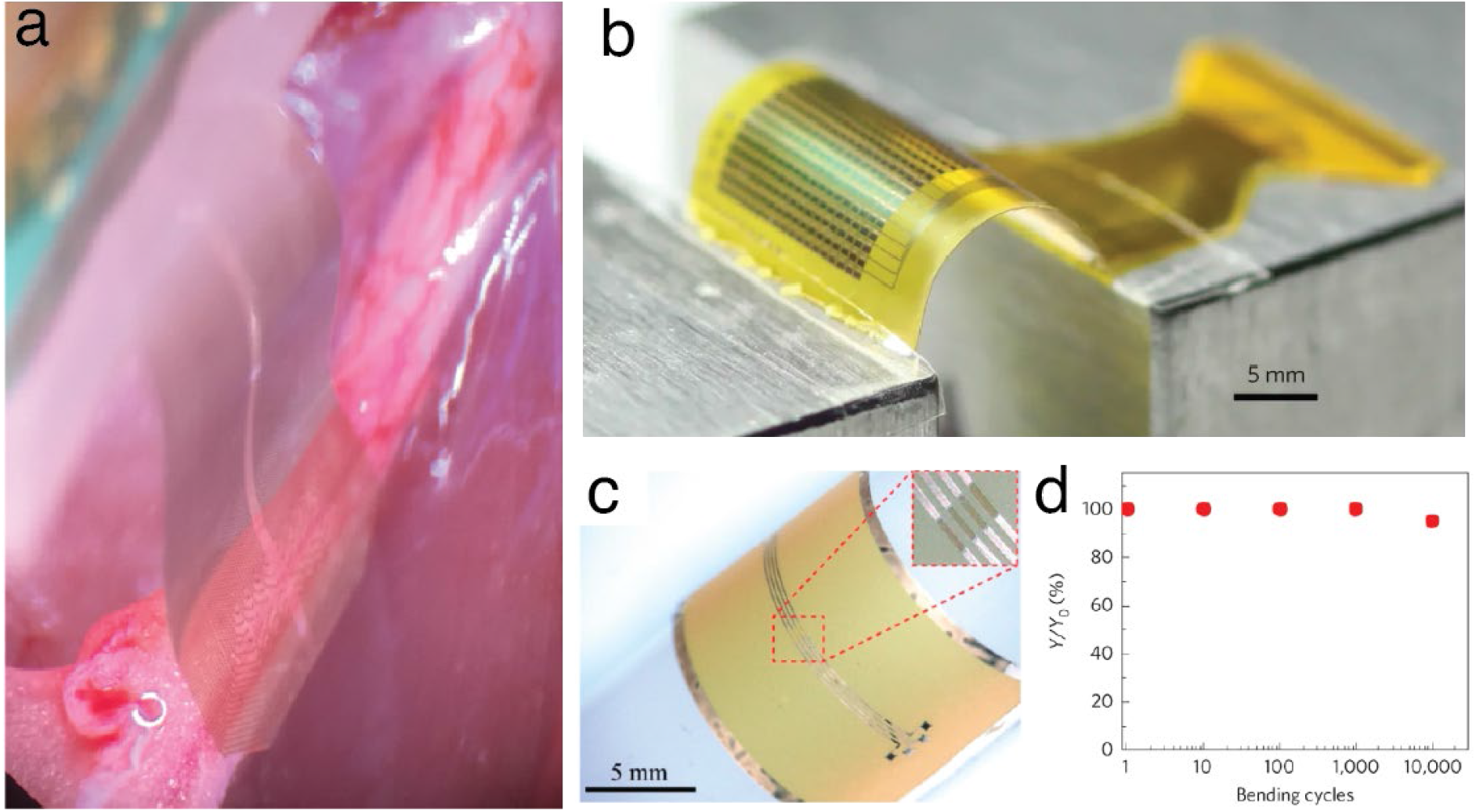
Flexible ceramic-encapsulated electronics enabled by thin films. a) SiC ECoG shown wrapped around the sciatic nerve of a rat (Diaz-Botia, et al., 2017). b) Thermal oxide encapsulated electrode array for cardiac monitoring (Fang, et al., 2016). c) SiC encapsulated electronics wrapped around a curved surface (radii of curvature 6 mm) (Phan, et al., 2019). d) percentage of working electrodes after a given number of bending cycles (bent to a radius of curvature of 5 mm) showing that the electronics are not compromised after bending cycles (Fang, et al., 2016).

At these thicknesses, however, degradation mechanisms such as hydrolysis begin to have non-negligible etch rates. For example, ALD Al_2_O_3_ is known to dissolve in water, hence the need to protect Al_2_O_3_ layers with CVD parylene or ALD titania (Abdulagatov, et al., 2011, Kim, et al., 2014). Furthermore, the material itself is not the only deciding factor for long-term stability of a thin film. The deposition method and parameters are critical to control film quality, bond-composition, morphology, among other physical properties. For example, thermal oxide films were shown by Fang et al. to have a dissolution rate in physiological conditions of roughly 0.04 nm/day, whereas a previous study by Maloney et al. showed a dissolution rate of 3.5 nm/day for PECVD silicon dioxide (Fang, et al., 2016, Maloney, et al., 2005). Similarly, Hsu et al. found that PECVD a-SiC films deposited at temperatures lower than 200 °C with low hydrogen dilution and high silane-to-methane precursor flow were dramatically less stable than a-SiC films deposited at higher temperatures in silane-starved conditions, which showed nearly no degradation in saline soak tests (Hsu, et al., 2007). Hermeticity testing of thin films is always essential; even small process variations can result in significant changes in film stability.

### 3.3 Quality testing for thin films

Several methods exist for leak testing thin films. He-leak testing can be performed but may be challenging due to the small sizes of the samples. One common method is to deposit the film-under-test over a test material which is affected by moisture. For example, magnesium (Mg) is highly reactive when exposed to water, hydrolyzing rapidly; a simple method for leak test begins by depositing a thin film of the material to be tested over a square of Mg, then creating a channel over the film and filling the channel with solution (Fig. 9a) (Fang, et al., 2016, Song, et al., 2017, Phan, et al., 2019). Any pinholes in the film or impurities which can be etched away act as become leakage paths for water to ingress through the film and eventually cause the underlying Mg film to react. The film can be inspected periodically with a microscope for evidence of etching or reaction under the thin film under test. In this way, time to failure can also be recorded and different films can be compared. Optical methods can be unreliable, however; some defects may be too small to identify. A different set of methods rely on shifts in material dielectric properties due to moisture ingress. As an example, resonant inductor-capacitor circuits can be fabricated from thin films or discrete components, thin-film coated with the material to be tested, then placed in a humid environment. By appropriate design of the inductor and proper tuning of the resonant circuit, the resonator can be wirelessly interrogated and impedance spectra shifts caused by changes in the components, putatively caused by moisture ingress (Fig. 9b) (Yeon, et al., 2019).

**Fig. 9.**
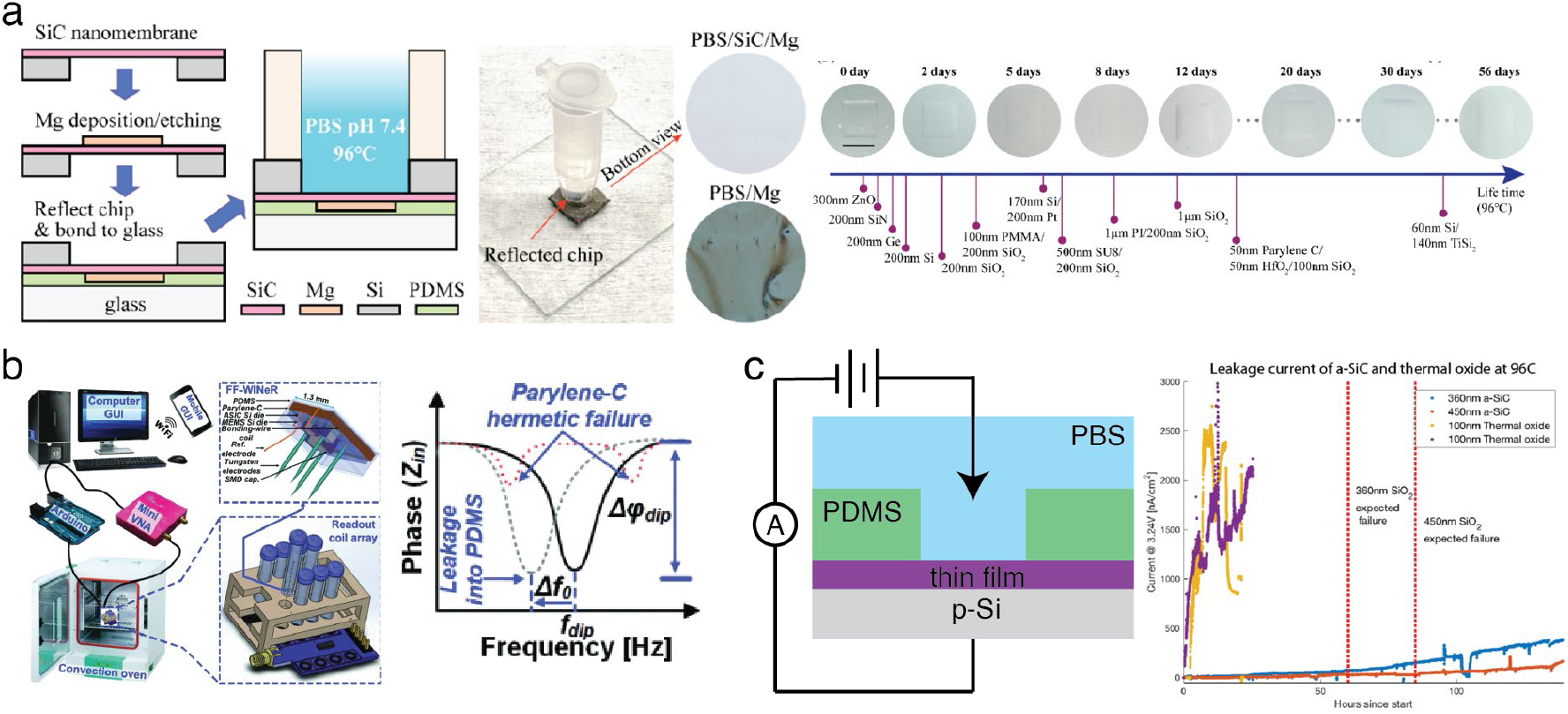
Methods of testing encapsulation hermiticity for miniaturized probes. a) Mg leak tests. Left: schematic of test set up. An SiC membrane is made, followed by the deposition of a square of Mg. The Mg layer is then potted in PDMS and plasma bonded to a glass slide for optical transparency. A Teflon tube filled with PBS is then placed over the Si emboss and the film is monitored for signs of Mg hydrolysis. Middle: photo of set up. Right: Timeline of thin films showing failure mode using this method. SiC shows the best performance (Phan, et al., 2019). b) Wireless testing method. Left: schematic of set up. Coated wireless microelectrode arrays are placed in a humid convection oven. A VNA controlled by Arduino measures the impedance spectrum of the system to determine leakage. Right: Mock figure showing how the phase spectrum shifts due to moisture ingress (Yeon, et al., 2019). c) Leakage current test. Left: schematic of set up. The thin film to test in question is deposited on a conductive substrate. A PDMS reservoir is created and placed over the thin film, then filled with PBS. A bias between the PBS and conductive substrate is applied and the leakage current is measured. Right: Figure showing leakage current as a function of time for a-SiC and thermal SiO_2_. a-SiC is shown to be a superior moisture barrier to thermal SiO_2_ (Diaz-Botia, et al., 2017).

Lastly, for materials acting as feedthrough or electrical insulators, any test should take into account that electrical potentials will exist across the encapsulation layer during operation. Electrical testing methods should apply these potentials during testing to accurately reproduce the implant environment. In leakage tests, this is usually done by applying a potential and measuring the resulting leakage current (Fig. 9c). Failure is determined when the leakage current rises above a set threshold current (Fang, et al., 2016, Diaz-Botia, et al., 2017, Haemmerli, et al., 2013).

## 4. Future Directions and Conclusion

There is no doubt that both bulk and thin-film ceramics will play a critical role in neural implants in contemporary and future devices. The use of alumina feedthroughs will remain an essential component of non-miniaturized AIMDs, as they currently are in cochlear implants, deep-brain-stimulation systems, and more. Microfabricated implants will also benefit from ceramic materials as the search for a thin-film hermetic encapsulation material continues. In this final section, we will examine some unique directions for implants that ceramic materials can offer.

### 4.1 Silicon carbide for encapsulating neural implants

SiC has emerged as one of the main contenders for chronic hermetic encapsulation due to its chemical stability, biocompatibility, and molecular barrier properties. The silicon-carbon bond in SiC is extremely stable: silicon atoms are bound to carbon atoms throughout the material, giving SiC its high chemical resistance (Maboudian, et al., 2013). As a result, proper tuning of deposition parameters can result in SiC thin films that exhibit little-to-no hydrolysis in-vivo or in-vitro (Cogan, et al., 2003, Lei, et al., 2016, Hsu, et al., 2007, Phan, et al., 2019). Properly tuned films have also been demonstrated to be excellent molecular barriers. Diaz-Botia et al. showed that 100 nm of a-SiC outperformed 100 nm thermal-SiO_2_ as a barrier to water vapor (Diaz-Botia, et al., 2016). Phan et al. showed that crystalline SiC exhibited very low ion diffusivity: after 12 weeks of soaking in 96 °C phosphate buffered saline (PBS), secondary ion mass spectroscopy revealed that by 100 nm through the film, no Na^+^ could be detected (Phan, et al., 2019). Additionally, the biocompatibility of SiC has been demonstrated both in-vivo and in-vitro (Cogan, et al., 2003, Saddow, et al., 2011, Frewin, et al., 2016). Cells cultured on SiC show better adhesion, proliferation, viability, and mitochondrial health than cells cultured on Si (Bonaventura, et al., 2019, Coletti, et al., 2007). SiC implants were also shown to have a decreased immune response compared to Si implants when utilized in-vivo. Frewin et al. found reduced microglia and macrophage activation attached to implanted SiC samples in rat tissue (Frewin, et al., 2011); Knaack et al. did not find significant differences between microglia and macrophage activation in a-SiC coated Si samples and uncoated Si samples, but did find reduced reactive astrocytes in a-SiC coated Si samples compared to the uncoated samples (Knaack, et al., 2016). Thus, while it is unclear if SiC is definitively more biocompatible than Si, it is evident that SiC is well tolerated by tissue and should be pursued as an encapsulation material. The Cogan group at the University of Texas at Dallas has demonstrated a-SiC encapsulated ultramicroelectrode planar arrays using and Utah electrode arrays (Deku, et al., 2018, Joshi-Imre, et al., 2019). Furthermore, because SiC is a semiconductor, it can be doped to become electrically conducting. Applications of electrically conductive SiC will be explored in the next section.

### 4.2 Seamless ceramic interfaces

It is well known that many packaging failure modes occur at interfaces of dissimilar materials; though Joshi-Imre et al. demonstrated that a-SiC encapsulated Utah electrode arrays did not show evidence of dissolution or cracking in the encapsulation layer (in contrast to parylene), delamination of the a-SiC was one failure mode of the implanted arrays (Joshi-Imre, et al., 2019). Unfortunately, heterogenous interfaces are unavoidable if feedthroughs are required. This failure point has been long recognized. For example, horizontal interconnects with vertical feedthroughs have been used to try to enhance hermiticity by increasing the length of the leakage path (Fig. 4b) (Guenther, et al., 2014). However, as conductor density increases within feedthroughs, ensuring hermiticity becomes more difficult as a single leak will result in a compromised device. Furthermore, miniaturization of devices will inevitably result in shorter leakage paths, increasing the risk of water vapor ingress.

Ceramic materials can provide a unique solution to this issue, in that some ceramics can be doped to vary the material from insulator to conductor. This raises the possibility that a feedthrough can be made of the same material, doped differently. Both penetrating and surface neural probes have been fabricated in all-SiC processes. Diaz-Botia et al.’s thin film SiC ECoGs utilized amorphous SiC as both the substrate and the encapsulation layer with n-SiC recording sites (Fig. 10a) and demonstrated electrical recordings of compound action potentials in the rat sciatic nerve (Diaz-Botia, et al., 2017). While complete device aging was not demonstrated in that work, the a-SiC thin film itself was aged in a benchtop reactive accelerated aging test and shown to be a superior encapsulation material to thermally grown SiO2. The Saddow group at the University of Southern Florida has also been exploring all-SiC neural probes using crystalline SiC (Bernardin, et al., 2018, Beygi, et al., 2019). In that body of work, n-type crystalline SiC substrates were used and p-type crystalline SiC interconnects were created to form reverse-biased diodes between interconnects for electrical isolation; a-SiC layer was used for encapsulation. While *in-vivo* results have not yet been demonstrated with these devices, the use of a homogenous material for all parts of the implant should greatly improve reliability.

**Fig. 10.**
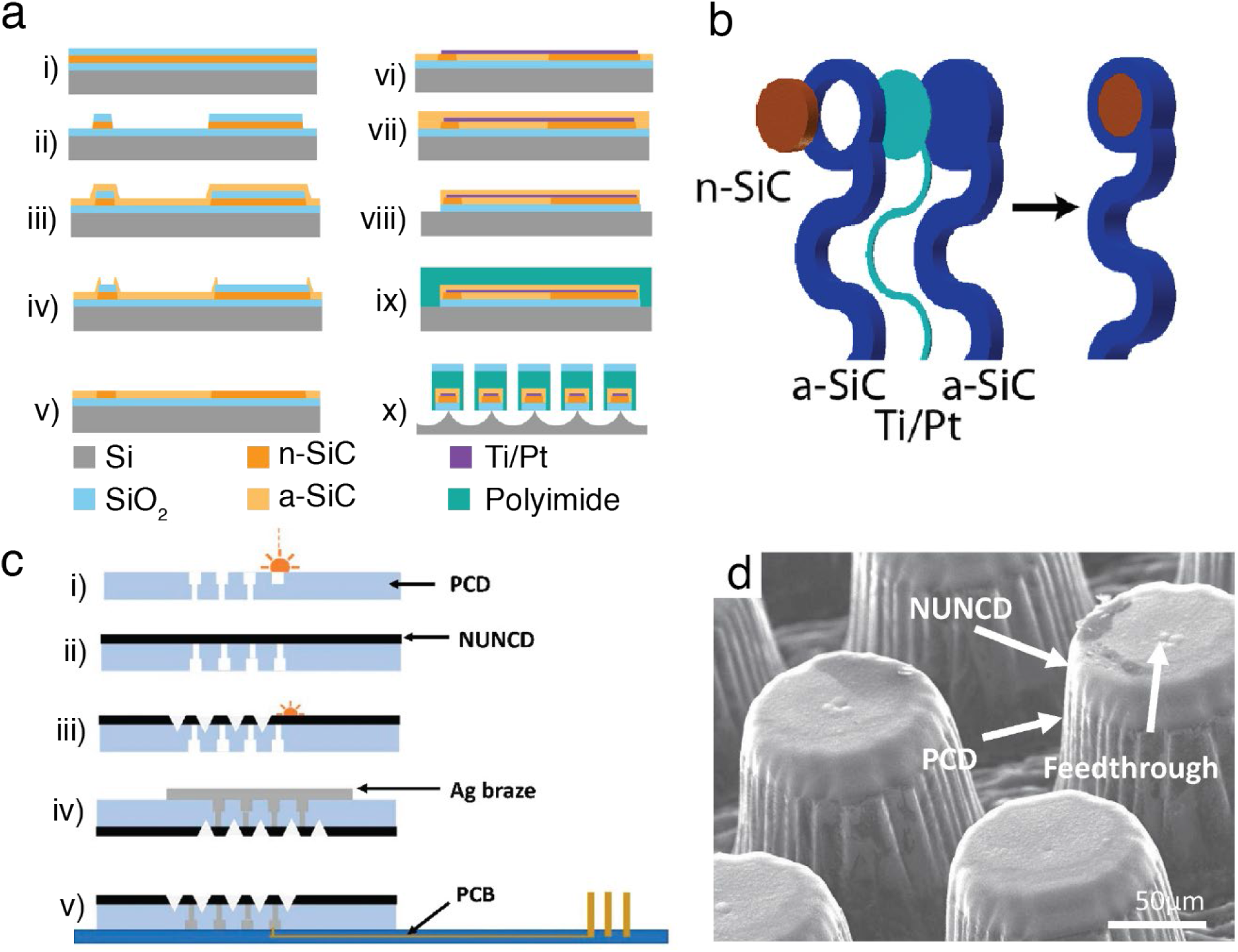
Process flows of seamless neural interfaces. a) Process flow of all-SiC ECoGs. i) an oxide layer is grown over a silicon substrate, n-doped SiC is deposited afterwards, and a second oxide layer is deposited on top. ii) the n-doped SiC and top oxide are then patterned to create electrodes and protect the electrodes from the next step, respectively. iii) a-SiC is then deposited to form the substrate a-SiC. iv) the first layer of a-SiC is then patterned, with the oxide layer acting as an etch stop. v) the top oxide is then stripped away and the free standing a-SiC is removed through sonication. vi) Ti/Pt interconnects are evaporated and patterned. vii) the encapsulating a-SiC layer is then deposited to fully encapsulate all components within a-SiC. viii) the a-SiC and underlying oxide are then patterned to form the device shape to reveal the final device. ix) prior to release, a polyimide backing layer is placed over the device to aid release and finally x) the device is released in XeF2. b) A 3D rendering of the electrode layers for the SiC ECoG in a) (Diaz-Botia, et al., 2017). c) Process flow for seamless diamond feedthroughs. i) a PCD substrate is laser milled to form vias. ii) a layer of N-UNCD is then grown over the substrate, which also covers the feedthroughs. iii) the N-UNCD is then patterned, again through laser milling. iv)to make connections to an external PCB, silver-ABA is applied over the substrate to fill the vias and fired. v) finally, the excess braze alloy is ground away through mechanical polishing, and the feedthrough array is flipchip bonded onto a PCB. d) SEM of the all diamond-feedthrough, showing the N-UNCD and PCD interface (Stamp, et al., 2020).

Diamond is another ceramic material that can exist in both insulating and conductive forms as PCD and nitrogen-doped ultra nano-crystalline diamond (N-UNCD), respectively. Ganesan et al. have utilized this property to create all-diamond feedthrough arrays for retinal prostheses by starting with a PCD substrate and then growing N-UNCD electrodes on top of it (Fig. 10b) (Ganesan, et al., 2014, Stamp, et al., 2020). These feedthroughs have been demonstrated as hermetic and biocompatible. However, the chemical inertness of diamond makes it very difficult to etch, and laser machining was used to pattern both the PCD and the N-UNCD, making scalability difficult. Nevertheless, there is much promise in the general principle of seamless neural interfaces, enabled by ceramic materials.

### 4.3 Ceramics for Acoustics

Ceramics have been a major enabler of electromagnetic wireless power transfer but they also play an important role in ultrasonic power transfer. The use of ultrasound in medical implants has been gaining popularity due to the efficiency of ultrasonic propagation in tissue compared to electromagnetics (Seo, et al., 2015, Seo, et al., 2016, Piech, et al., 2017, Piech, et al., 2020, Ghanbari, et al., 2019, Vo, et al., 2020, Charthad, et al., 2018, Weber, et al., 2018, Chang, et al., 2017, Arbabian, et al., 2016, Charthad, et al., 2015, Sonmezoglu et al., 2020). Functional piezoelectric ceramics, such as lead-zirconate-titanate (PZT) and barium titanate (BaTiO_3_), have been used extensively as power harvesting transducers to convert incoming mechanical energy to electrical energy thereby powering implant microelectronics. While the use of ultrasound allows for metallic housings (Shen & Maharbiz, 2019), engineered ceramics with nanoscale grain-size and low porosity have been shown to be good acoustic windows in non-active medical implants for ultrasound-based therapies (Gutierrez, et al., 2017). Thus, ceramic housings not only enable visible-to-RF EM transparency, but also can enable acoustic transparency.

### 4.4 Conclusion

Ceramic materials have great potential in the packaging of neural implants due to their biocompatibility, corrosion resistance, RF transparency, and tunable dielectric properties. Although bulk ceramic can be difficult to shape and can be prone to cracking, thin-film ceramics are not as susceptible to these issues. Metals and polymers will continue to have a central place in the packaging of neural probes but ceramics provide a complementary pathway to reliable, aggressively miniaturized, high electrode density, wireless AIMDs.

## References

1. Abdulagatov, A. I. et al., 2011. Al2O3 and TiO2 atomic layer deposition on copper for water corrosion resistance. ACS applied materials & interfaces, Volume 3, pp. 4593–4601.

2. Agathopoulos, S., Correia, R. N., Joanni, E. & Fernandes, J. R. A., 2002. Interactions at zirconia-Au-Ti interfaces at high temperatures.

3. Allen, R. V. & Borbidge, W. E., 1983. Solid state metal-ceramic bonding of platinum to alumina. Journal of Materials Science, Volume 18, pp. 2835–2843.

4. Apollo, N. V. et al., 2016. Brazing techniques for the fabrication of biocompatible carbon-based electronic devices. Carbon, Volume 107, pp. 180–189.

5. Arbabian, A. et al., 2016. Sound technologies, sound bodies: Medical implants with ultrasonic links. IEEE Microwave Magazine, Volume 17, pp. 39–54.

6. Barrese, J. C. et al., 2013. Failure mode analysis of silicon-based intracortical microelectrode arrays in non-human primates. Journal of neural engineering, Volume 10, p. 066014.

7. Bernardin, E. K. et al., 2018. Demonstration of a robust all-silicon-carbide intracortical neural interface. Micromachines, Volume 9, p. 412.

8. Beygi, M. et al., 2019. Fabrication of a Monolithic Implantable Neural Interface from Cubic Silicon Carbide. Micromachines, Volume 10, p. 430.

9. Bjune, C. K. et al., 2015. Package architecture and component design for an implanted neural stimulator with closed loop control. s.l., s.n., pp. 7825–7830.

10. Bonaventura, G. et al., 2019. Biocompatibility between Silicon or Silicon carbide surface and neural Stem cells. Scientific reports, Volume 9, pp. 1–13.

11. Borovikova, L. V. et al., 2000. Vagus nerve stimulation attenuates the systemic inflammatory response to endotoxin. Nature, Volume 405, pp. 458–462.

12. Borton, D. A., Yin, M., Aceros, J. & Nurmikko, A., 2013. An implantable wireless neural interface for recording cortical circuit dynamics in moving primates. Journal of neural engineering, Volume 10, p. 026010.

13. Bosch, J. L. H. R. & Groen, J., 2000. Sacral nerve neuromodulation in the treatment of patients with refractory motor urge incontinence: long-term results of a prospective longitudinal study. The Journal of urology, Volume 163, pp. 1219–1222.

14. Bowman, L. & Meindl, J. D., 1986. The packaging of implantable integrated sensors. IEEE Transactions on Biomedical Engineering, pp. 248–255.

15. Buzsáki, G., 2004. Large-scale recording of neuronal ensembles. Nature neuroscience, Volume 7, pp. 446–451.

16. Cavuoto, J., 2018. The market for neurotechnology: 2018--2022. Neurotech Report, Volume 1.

17. Cha, K., Horch, K. & Normann, R. A., 1992. Simulation of a phosphene-based visual field: visual acuity in a pixelized vision system. Annals of biomedical engineering, Volume 20, pp. 439–449.

18. Chang, E. F., 2015. Towards large-scale, human-based, mesoscopic neurotechnologies. Neuron, Volume 86, pp. 68–78.

19. Chang, T. C. et al., 2017. Scaling of ultrasound-powered receivers for sub-millimeter wireless implants. s.l., s.n., pp. 1–4.

20. Charthad, J. et al., 2018. A mm-sized wireless implantable device for electrical stimulation of peripheral nerves. IEEE transactions on biomedical circuits and systems, Volume 12, pp. 257–270.

21. Charthad, J., Weber, M. J., Chang, T. C. & Arbabian, A., 2015. A mm-sized implantable medical device (IMD) with ultrasonic power transfer and a hybrid bi-directional data link. IEEE Journal of solid-state circuits, Volume 50, pp. 1741–1753.

22. Chen, R., Canales, A. & Anikeeva, P., 2017. Neural recording and modulation technologies. Nature Reviews Materials, Volume 2, pp. 1–16.

23. Chen, Z. et al., 2019. 3D printing of ceramics: A review. Journal of the European Ceramic Society, Volume 39, pp. 661–687.

24. Cheung, K. C., 2007. Implantable microscale neural interfaces. Biomedical microdevices, Volume 9, pp. 923–938.

25. Christensen, M. B. et al., 2014. The foreign body response to the Utah Slant Electrode Array in the cat sciatic nerve. Acta biomaterialia, Volume 10, pp. 4650–4660.

26. Christian & Kenis, P. J. A., 2007. Fabrication of ceramic microscale structures. Journal of the American Ceramic Society, Volume 90, pp. 2779–2783.

27. Chung, J. E. et al., 2019. High-density, long-lasting, and multi-region electrophysiological recordings using polymer electrode arrays. Neuron, Volume 101, pp. 21–31.

28. Cogan, S. F. et al., 2003. Plasma-enhanced chemical vapor deposited silicon carbide as an implantable dielectric coating. Journal of Biomedical Materials Research Part A: An Official Journal of The Society for Biomaterials, The Japanese Society for Biomaterials, and The Australian Society for Biomaterials and the Korean Society for Biomaterials, Volume 67, pp. 856–867.

29. Coletti, C. et al., 2007. Biocompatibility and wettability of crystalline SiC and Si surfaces. s.l., s.n., pp. 5849–5852.

30. Correia, R. N., Emiliano, J. V. & Moretto, P., 1998. Microstructure of diffusional zirconia--titanium and zirconia--(Ti--6 wt% Al--4 wt% V) alloy joints. Journal of materials science, Volume 33, pp. 215–221.

31. De Paris, A., Robin, M. & Fantozzi, G., 1991. Welding of ceramics SiO2-Al2O3 by laser beam. Le Journal de Physique IV, Volume 1, pp. C7--127.

32. Deku, F. et al., 2018. Amorphous silicon carbide ultramicroelectrode arrays for neural stimulation and recording. Journal of neural engineering, Volume 15, p. 016007.

33. Diaz-Botia, C. A. et al., 2017. A silicon carbide array for electrocorticography and peripheral nerve recording. Journal of Neural Engineering, Volume 14, p. 056006.

34. Donaldson, P. E. K., 1976. The encapsulation of microelectronic devices for long-term surgical implantation. IEEE Transactions on Biomedical Engineering, pp. 281–285.

35. Donaldson, P. E. K. & Sayer, E., 1981. A technology for implantable hermetic packages. Part 1: Design and materials. Medical and Biological Engineering and Computing, Volume 19, p. 398.

36. Elterman, D. S., 2018. The novel Axonics^®^ rechargeable sacral neuromodulation system: Procedural and technical impressions from an initial North American experience. Neurourology and urodynamics, Volume 37, pp. S1––S8.

37. Exner, H. & Nagel, A.-M., 1999. Laser welding of functional and constructional ceramics for Microelectronics. s.l., s.n., pp. 262–268.

38. Famm, K. et al., 2013. Drug discovery: a jump-start for electroceuticals. Nature, Volume 496, p. 159.

39. Fang, H. et al., 2017. Capacitively coupled arrays of multiplexed flexible silicon transistors for long-term cardiac electrophysiology. Nature biomedical engineering, Volume 1, pp. 1–12.

40. Fang, H. et al., 2016. Ultrathin, transferred layers of thermally grown silicon dioxide as biofluid barriers for biointegrated flexible electronic systems. Proceedings of the National Academy of Sciences, Volume 113, pp. 11682–11687.

41. Frewin, C. L. et al., 2016. Silicon Carbide as a Robust Neural Interface. ECS Transactions, Volume 75, p. 39.

42. Frewin, C. L., Locke, C., Saddow, S. E. & Weeber, E. J., 2011. Single-crystal cubic silicon carbide: An in vivo biocompatible semiconductor for brain machine interface devices. s.l., s.n., pp. 2957–2960.

43. Ganesan, K. et al., 2014. An all-diamond, hermetic electrical feedthrough array for a retinal prosthesis. Biomaterials, Volume 35, pp. 908–915.

44. Gedopt, J. & Delarbre, E., 2000. Pulsed Nd: YAG laser welding of titanium ear implants. s.l., s.n., pp. 264–267.

45. Ghanbari, M. M. et al., 2019. A Sub-mm 3 Ultrasonic Free-Floating Implant for Multi-Mote Neural Recording. IEEE Journal of Solid-State Circuits, Volume 54, pp. 3017–3030.

46. Gilbert, J. L., Buckley, C. A. & Jacobs, J. J., 1993. In vivo corrosion of modular hip prosthesis components in mixed and similar metal combinations. The effect of crevice, stress, motion, and alloy coupling. Journal of biomedical materials research, Volume 27, pp. 1533–1544.

47. Gill, E. C. et al., 2013. High-density feedthrough technology for hermetic biomedical micropackaging. MRS Online Proceedings Library Archive, Volume 1572.

48. Greenhouse, H., 2000. Hermeticity of electronic packages. s.l.:Elsevier.

49. Green, R. A. et al., 2013. Integrated electrode and high density feedthrough system for chip-scale implantable devices. Biomaterials, Volume 34, pp. 6109–6118.

50. Grill, W. M. & Mortimer, J. T., 2000. Neural and connective tissue response to long-term implantation of multiple contact nerve cuff electrodes. Journal of Biomedical Materials Research: An Official Journal of The Society for Biomaterials, The Japanese Society for Biomaterials, and The Australian Society for Biomaterials and the Korean Society for Biomaterials, Volume 50, pp. 215–226.

51. Guenther, T. et al., 2014. Pt-Al2O3 interfaces in cofired ceramics for use in miniaturized neuroprosthetic implants. Journal of Biomedical Materials Research Part B: Applied Biomaterials, Volume 102, pp. 500–507.

52. Guenther, T. et al., 2020. Practical aspects and limitations of hermeticity testing of micro-encapsulations using cumulative helium leak detection for miniaturized implantable medical devices. IEEE Transactions on Components, Packaging and Manufacturing Technology.

53. Gutierrez, M. I. et al., 2017. Novel cranial implants of yttria-stabilized zirconia as acoustic windows for ultrasonic brain therapy. Advanced healthcare materials, Volume 6, p. 1700214.

54. Haemmerli, A. J. et al., 2013. Ultra-thin atomic layer deposition films for corrosion resistance. s.l., s.n., pp. 1931–1934.

55. Hsu, J.-M.et al., 2007. Characterization of a-SiCx: H thin films as an encapsulation material for integrated silicon based neural interface devices. Thin solid films, Volume 516, pp. 34–41.

56. James, R. J., Gobet, J., Durante, G. S. & Fretz, M., 2016. A novel packaging technology for miniaturization of active long-term implantable medical devices enabling new medical applications. s.l., s.n., pp. 1–6.

57. Jeong, J. et al., 2019. Conformal hermetic sealing of wireless microelectronic implantable chiplets by multilayered atomic layer deposition (ALD). Advanced Functional Materials, Volume 29, p. 1806440.

58. Jiang, G. et al., 2005. Zirconia to Ti-6Al-4V braze joint for implantable biomedical device. Journal of Biomedical Materials Research Part B: Applied Biomaterials: An Official Journal of The Society for Biomaterials, The Japanese Society for Biomaterials, and The Australian Society for Biomaterials and the Korean Society for Biomaterials, Volume 72, pp. 316–321.

59. Jiang, G. & Zhou, D. D., 2009. Technology advances and challenges in hermetic packaging for implantable medical devices. In: Implantable Neural Prostheses 2. s.l.:Springer, pp. 27–61.

60. Joshi-Imre, A. et al., 2019. Chronic recording and electrochemical performance of amorphous silicon carbide-coated Utah electrode arrays implanted in rat motor cortex. Journal of neural engineering, Volume 16, p. 046006.

61. Jun, J. J. et al., 2017. Fully integrated silicon probes for high-density recording of neural activity. Nature, Volume 551, pp. 232–236.

62. Kane, M. J., Breen, P. P., Quondamatteo, F. & ÓLaighin, G., 2011. BION microstimulators: A case study in the engineering of an electronic implantable medical device. Medical engineering & physics, Volume 33, pp. 7–16.

63. Kelly, S. K. et al., 2011. A hermetic wireless subretinal neurostimulator for vision prostheses. IEEE transactions on biomedical engineering, Volume 58, pp. 3197–3205.

64. Kennell, G. F. & Evitts, R. W., 2009. Crevice corrosion cathodic reactions and crevice scaling laws. Electrochimica Acta, Volume 54, pp. 4696–4703.

65. Kim, L. H. et al., 2014. Al2O3/TiO2 nanolaminate thin film encapsulation for organic thin film transistors via plasma-enhanced atomic layer deposition. ACS applied materials & interfaces, Volume 6, pp. 6731–6738.

66. Knaack, G. L. et al., 2016. In vivo characterization of amorphous silicon carbide as a biomaterial for chronic neural interfaces. Frontiers in neuroscience, Volume 10, p. 301.

67. Kozai, T. D. Y. et al., 2015. Brain tissue responses to neural implants impact signal sensitivity and intervention strategies. ACS chemical neuroscience, Volume 6, pp. 48–67.

68. Langenmair, M. et al., 2018. Low temperature approach for high density electrical feedthroughs for neural implants using maskless fabrication techniques. s.l., s.n., pp. 2933–2936.

69. Lei, X. et al., 2016. SiC protective coating for photovoltaic retinal prosthesis. Journal of neural engineering, Volume 13, p. 046016.

70. Lichter, S. G. et al., 2015. Hermetic diamond capsules for biomedical implants enabled by gold active braze alloys. Biomaterials, Volume 53, pp. 464–474.

71. Lind, G., Linsmeier, C. E. & Schouenborg, J., 2013. The density difference between tissue and neural probes is a key factor for glial scarring. Scientific reports, Volume 3, p. 2942.

72. Li, W., Kabius, B. & Auciello, O., 2010. Science and technology of biocompatible thin films for implantable biomedical devices. s.l., s.n., pp. 6237–6242.

73. Loeb, G. E. et al., 2007. Mechanical loading of rigid intramuscular implants. Biomedical microdevices, Volume 9, pp. 901–910.

74. Lovell, N. H. et al., 2005. A retinal neuroprosthesis design based on simultaneous current injection. s.l., s.n., pp. 98–101.

75. Maboudian, R., Carraro, C., Senesky, D. G. & Roper, C. S., 2013. Advances in silicon carbide science and technology at the micro-and nanoscales. Journal of Vacuum Science & Technology A: Vacuum, Surfaces, and Films, Volume 31, p. 050805.

76. Maloney, J. M., Lipka, S. A. & Baldwin, S. P., 2005. In vivo biostability of CVD silicon oxide and silicon nitride films. MRS Online Proceedings Library Archive, Volume 872.

77. Massey, T. L. et al., 2019. A high-density carbon fiber neural recording array technology. Journal of neural engineering, Volume 16, p. 016024.

78. Miyamoto, I., Cvecek, K., Okamoto, Y. & Schmidt, M., 2014. Internal modification of glass by ultrashort laser pulse and its application to microwelding. Applied Physics A, Volume 114, pp. 187–208.

79. Najafi, K., Wise, K. D. & Mochizuki, T., 1985. A high-yield IC-compatible multichannel recording array. IEEE Transactions on Electron Devices, Volume 32, pp. 1206–1211.

80. Normann, R. A. & Fernandez, E., 2016. Clinical applications of penetrating neural interfaces and Utah Electrode Array technologies. Journal of neural engineering, Volume 13, p. 061003.

81. Patrick, E., Orazem, M. E., Sanchez, J. C. & Nishida, T., 2011. Corrosion of tungsten microelectrodes used in neural recording applications. Journal of neuroscience methods, Volume 198, pp. 158–171.

82. Peña, E. et al., 2008. Skin erosion over implants in deep brain stimulation patients. Stereotactic and functional neurosurgery, Volume 86, pp. 120–126.

83. Penilla, E. H. et al., 2019. Ultrafast laser welding of ceramics. Science, Volume 365, pp. 803–808.

84. Phan, H.-P.et al., 2019. Long-Lived, Transferred Crystalline Silicon Carbide Nanomembranes for Implantable Flexible Electronics. ACS nano, Volume 13, pp. 11572–11581.

85. Piconi, C. & Maccauro, G., 1999. Zirconia as a ceramic biomaterial. Biomaterials, Volume 20, pp. 1–25.

86. Piech, D. K. et al., 2020. A wireless millimetre-scale implantable neural stimulator with ultrasonically powered bidirectional communication. Nature Biomedical Engineering, Volume 4, pp. 207–222.

87. Piech, D. K., Kay, J. E., Boser, B. E. & Maharbiz, M. M., 2017. Rodent wearable ultrasound system for wireless neural recording. s.l., s.n., pp. 221–225.

88. Polikov, V. S., Tresco, P. A. & Reichert, W. M., 2005. Response of brain tissue to chronically implanted neural electrodes. Journal of neuroscience methods, Volume 148, pp. 1–18.

89. Poon, A. S. Y., 2014. Miniaturized Biomedical Implantable Devices1. Edited by Evgeny Katz.

90. Potter, K. A., Buck, A. C., Self, W. K. & Capadona, J. R., 2012. Stab injury and device implantation within the brain results in inversely multiphasic neuroinflammatory and neurodegenerative responses. Journal of neural engineering, Volume 9, p. 046020.

91. Raducanu, B. C. et al., 2017. Time multiplexed active neural probe with 1356 parallel recording sites. Sensors, Volume 17, p. 2388.

92. Riviere, C., Robin, M. & Fantozzi, G., 1994. Comparison between two techniques in laser welding of ceramics. Le Journal de Physique IV, Volume 4, pp. C4--135.

93. Romero, E. et al., 2001. Neural morphological effects of long-term implantation of the self-sizing spiral cuff nerve electrode. Medical and Biological Engineering and Computing, Volume 39, pp. 90–100.

94. Rouse, A. G. et al., 2011. A chronic generalized bi-directional brain--machine interface. Journal of neural engineering, Volume 8, p. 036018.

95. Ruys, A. J., 2018. Alumina Ceramics: Biomedical and Clinical Applications. s.l.:Woodhead Publishing.

96. Saddow, S. E. et al., 2011. Single-crystal silicon carbide: A biocompatible and hemocompatible semiconductor for advanced biomedical applications. s.l., s.n., pp. 824–830.

97. Salatino, J. W., Ludwig, K. A., Kozai, T. D. Y. & Purcell, E. K., 2017. Glial responses to implanted electrodes in the brain. Nature biomedical engineering, Volume 1, pp. 862–877.

98. Santella, M. L. & Pak, J. J., 1993. Brazing titanium-vapor-coated zirconia. WELDING JOURNAL-NEW YORK-, Volume 72, pp. 165--s.

99. Schuettler, M. et al., 2010. A device for vacuum drying, inert gas backfilling and solder sealing of hermetic implant packages. s.l., s.n., pp. 1577–1580.

100. Schuettler, M. et al., 2010. Fabrication and test of a hermetic miniature implant package with 360 electrical feedthroughs. s.l., s.n., pp. 1585–1588.

101. Schuettler, M. & Stieglitz, T., 2013. Microassembly and micropackaging of implantable systems. In: Implantable Sensor Systems for Medical Applications. s.l.:Elsevier, pp. 108–149.

102. Seo, D. et al., 2015. Model validation of untethered, ultrasonic neural dust motes for cortical recording. Journal of neuroscience methods, Volume 244, pp. 114–122.

103. Seo, D. et al., 2016. Wireless recording in the peripheral nervous system with ultrasonic neural dust. Neuron, Volume 91, pp. 529–539.

104. Seymour, J. P. & Kipke, D. R., 2007. Neural probe design for reduced tissue encapsulation in CNS. Biomaterials, Volume 28, pp. 3594–3607.

105. Seymour, J. P., Wu, F., Wise, K. D. & Yoon, E., 2017. State-of-the-art MEMS and microsystem tools for brain research. Microsystems & Nanoengineering, Volume 3, p. 16066.

106. Shah, K. G. et al., 2012. HIGH-DENSITY, BIO-COMPATIBLE, AND HERMETIC ELECTRICAL FEEDTHROUGHS USING EXTRUDED METAL VIAS, s.l.: s.n.

107. Shen, K. & Maharbiz, M. M., 2019. Design of Ceramic Packages for Ultrasonically Coupled Implantable Medical Devices. IEEE Transactions on Biomedical Engineering.

108. Siddiqui, M. S. & Jones, W. K., 2014. Vacuum Brazing of Alumina to Titanium for Implantable Feedthroughs Using Pure Gold as the Braze Metal. International Journal of Materials Science and Engineering, Volume 2, pp. 56–62.

109. Slavin, K. V., Nersesyan, H. & Wess, C., 2006. Peripheral neurostimulation for treatment of intractable occipital neuralgia. Neurosurgery, Volume 58, pp. 112–119.

110. Song, E. et al., 2017. Thin, transferred layers of silicon dioxide and silicon nitride as water and ion barriers for implantable flexible electronic systems. Advanced Electronic Materials, Volume 3, p. 1700077.

111. Song, E. et al., 2018. Transferred, ultrathin oxide bilayers as biofluid barriers for flexible electronic implants. Advanced Functional Materials, Volume 28, p. 1702284.

112. Stamp, M. E. M. et al., 2020. 3D Diamond Electrode Array for High-Acuity Stimulation in Neural Tissue. ACS Applied Bio Materials, Volume 3, pp. 1544–1552.

113. Steinmetz, N. A., Koch, C., Harris, K. D. & Carandini, M., 2018. Challenges and opportunities for large-scale electrophysiology with Neuropixels probes. Current opinion in neurobiology, Volume 50, pp. 92–100.

114. Stevenson, I. H. & Kording, K. P., 2011. How advances in neural recording affect data analysis. Nature neuroscience, Volume 14, p. 139.

115. Stieglitz, T., 2010. Manufacturing, assembling and packaging of miniaturized neural implants. Microsystem technologies, Volume 16, pp. 723–734.

116. Stöver, T. & Lenarz, T., 2009. Biomaterials in cochlear implants. GMS current topics in otorhinolaryngology, head and neck surgery, Volume 8.

117. Sun, F. T. & Morrell, M. J., 2014. Closed-loop neurostimulation: the clinical experience. Neurotherapeutics, Volume 11, pp. 553–563.

118. Szarowski, D. H. et al., 2003. Brain responses to micro-machined silicon devices. Brain research, Volume 983, pp. 23–35.

119. Takmakov, P. et al., 2015. Rapid evaluation of the durability of cortical neural implants using accelerated aging with reactive oxygen species. Journal of neural engineering, Volume 12, p. 026003.

120. Tracey, K. J., 2002. The inflammatory reflex. Nature, Volume 420, pp. 853–859.

121. Traeger, R., 1977. Nonhermeticity of polymeric lid sealants. IEEE Transactions on parts, Hybrids, and packaging, Volume 13, pp. 147–152.

122. Travitzky, N. et al., 2014. Additive manufacturing of ceramic-based materials. Advanced Engineering Materials, Volume 16, pp. 729–754.

123. Tummala, R. R., 1991. Ceramic and glass-ceramic packaging in the 1990s. Journal of the American Ceramic Society, Volume 74, pp. 895–908.

124. Turner, J. N. et al., 1999. Cerebral astrocyte response to micromachined silicon implants. Experimental neurology, Volume 156, pp. 33–49.

125. Vanhoestenberghe, A. & Donaldson, N., 2011. The limits of hermeticity test methods for micropackages. Artificial organs, Volume 35, pp. 242–244.

126. Vanhoestenberghe, A. & Donaldson, N., 2013. Corrosion of silicon integrated circuits and lifetime predictions in implantable electronic devices. Journal of neural engineering, Volume 10, p. 031002.

127. Vanhoestenberghe, A., Donaldson, N., Lovell, N. H. & Suaning, G. J., 2008. Hermetic encapsulation of an implantable vision prosthesis-Combining implant fabrication philosophies. s.l., s.n., pp. 187–9.

128. Viventi, J. et al., 2011. Flexible, foldable, actively multiplexed, high-density electrode array for mapping brain activity in vivo. Nature neuroscience, Volume 14, p. 1599.

129. Vlasov, A. S. & Karabanova, T. A., 1993. Ceramics and medicine. Glass and ceramics, Volume 50, pp. 398–401.

130. Vo, J. et al., 2020. Assessment of miniaturized ultrasound-powered implants: an in vivo study. Journal of Neural Engineering, Volume 17, p. 016072.

131. Watanabe, W. et al., 2006. Space-selective laser joining of dissimilar transparent materials using femtosecond laser pulses. Applied physics letters, Volume 89, p. 021106.

132. Weber, M. J. et al., 2018. A miniaturized single-transducer implantable pressure sensor with time-multiplexed ultrasonic data and power links. IEEE Journal of Solid-State Circuits, Volume 53, pp. 1089–1101.

133. Weiland, J. D. et al., 2013. Chip-scale packaging for bioelectronic implants. s.l., s.n., pp. 931–936.

134. Wellman, S. M. et al., 2019. Revealing spatial and temporal patterns of cell death, glial proliferation, and blood-brain barrier dysfunction around implanted intracortical neural interfaces. Frontiers in neuroscience, Volume 13, p. 493.

135. Wise, K. D., 2005. Silicon microsystems for neuroscience and neural prostheses. IEEE Engineering in Medicine and Biology Magazine, Volume 24, pp. 22–29.

136. Witte, R., Herfurth, H.-J. & Heinemann, S., 2002. Laser joining of glass with silicon. s.l., s.n., pp. 487–495.

137. Wu, Q., Lorenz, N., Cannon, K. M. & Hand, D. P., 2010. Glass frit as a hermetic joining layer in laser based joining of miniature devices. IEEE Transactions on components and packaging technologies, Volume 33, pp. 470–477.

138. Xiansheng, N., Zhenggan, Z., Xiongwei, W. & Luming, L., 2011. The use of Taguchi method to optimize the laser welding of sealing neuro-stimulator. Optics and Lasers in Engineering, Volume 49, pp. 297–304.

139. Xie, X. et al., 2013. Long-Term Bilayer Encapsulation Performance of Atomic Layer Deposited Al {bf 2} O {bf 3} and Parylene C for Biomedical Implantable Devices. IEEE Transactions on Biomedical Engineering, Volume 60, pp. 2943–2951.

140. Yang, H. et al., 2018. Chronically implantable package based on alumina ceramics and titanium with high-density feedthroughs for medical implants. s.l., s.n., pp. 3382–3385.

141. Yeong, W., Yap, C. Y., Mapar, M. & Chua, C. K., 2013. State-of-the-art review on selective laser melting of ceramics. High value manufacturing: advanced research in virtual and rapid prototyping, Volume 1, pp. 65–70.

142. Yeon, P. et al., 2019. Automated High-Throughput Hermetic Failure Monitoring System for Millimeter-Sized Wireless Implantable Medical Devices. s.l., s.n., pp. 2235–2238.

143. Zhou, A. et al., 2019. A wireless and artefact-free 128-channel neuromodulation device for closed-loop stimulation and recording in non-human primates. Nature biomedical engineering, Volume 3, pp. 15–26.

144. Zimmermann, F. et al., 2013. Ultrastable bonding of glass with femtosecond laser bursts. Applied Optics, Volume 52, pp. 1149–1154.

